# Localized nanoparticle-mediated delivery of miR-29b normalises the dysregulation of bone homeostasis caused by osteosarcoma whilst simultaneously inhibiting tumour growth

**DOI:** 10.1101/2022.09.09.507272

**Authors:** Fiona E. Freeman, Pere Dosta, Cristobal J. Riojas Javelly, Olwyn R. Mahon, Daniel J. Kelly, Natalie Artzi

## Abstract

Patients diagnosed with osteosarcoma undergo extensive surgical intervention and chemotherapy resulting in dismal prognosis and compromised quality of life owing to poor bone regeneration, which is further compromised with chemotherapy delivery. This study aims to investigate if localised delivery of miR-29b—which has been shown to promote bone formation by inducing osteoblast differentiation and also to suppress prostate and glioblastoma tumour growth—would suppress osteosarcoma tumours whilst simultaneously normalising the dysregulation of bone homeostasis caused by osteosarcoma. Thus, we studied the therapeutic potential of miR-29b to promote bone remodelling in an orthotopic model of osteosarcoma (rather than in bone defect models using healthy mice), and in the context of chemotherapy, that is clinically relevant. We developed a formulation of miR-29b:nanoparticles that were delivered via a novel hyaluronic-based hydrogel to enable local and sustained release of the therapy, and to study the potential of attenuating tumour growth whilst normalising bone homeostasis. We found that when miR-29b was delivered along with systemic chemotherapy, compared to chemotherapy alone, our therapy provided a significant decrease in tumour burden, increase in mouse survival, and a significant decrease in osteolysis thereby normalising the dysregulation of bone lysis activity caused by the tumour.

## Introduction

Osteosarcoma is a highly aggressive bone cancer, largely affecting children and adolescents with a worldwide incidence rate of 3,400 new cases per year[1]. Due to the young age of initial diagnosis, the management of this disease is a challenging and costly exercise, estimated to be €14.7 billion in Europe in the last 18 years[2]. Despite significant advances in treatment seen in other malignancies, no major improvements in outcome have been achieved since the introduction of chemotherapy in the 1970s[3]. Contemporary chemotherapy for osteosarcoma is normally a combination of one or two different chemotherapeutic drugs; doxorubicin or cisplatin, delivered to the patient either intravenously or orally. This chemotherapy is usually administered to the patient with 3 cycles prior to tumour resection and 3 cycles post tumour resection. However, these chemotherapeutic drugs have a series of short and long-term side effects on the patients and are not curative[4]. This emphasises the strong clinical unmet need for newer, more effective, treatment options to improve the overall survival of these young patients[5].

As osteosarcoma is such an aggressive disease, the surgical intervention usually involves total reconstruction of the limbs or in most cases amputation. To add to this, both chemotherapeutics and osteosarcoma tumours have been shown to disrupt bone homeostasis[6], resulting in a dysregulated bone lysis activity significantly hindering the surrounding bone’s ability to regenerate following surgical intervention. Therefore, any bone regeneration strategy that would stimulate the bone remodelling process, within the surrounding bone following surgical intervention, would be of great interest to these young patients. There exists a fine balance between trying to promote bone remodelling and promoting tumour growth, which has significantly slowed down fundamental research and clinical translation of tissue engineering strategies for cancer patients[7]. Clinically, the use of artificial bone graft substitutes only occurs following the removal of low-grade malignant bone tumours[8], and thus not in the case of osteosarcoma. While the use of growth factors like Bone Morphogenic Protein-2 (BMP-2) after malignant tumour resection remains a controversial topic. To understand this further, we previously developed a novel 3D co-culture spheroid model for early- and late-stage osteosarcoma[7]. Using this model, we demonstrated that BMP-2 delivery had a pro-osteogenic effect on the surrounding stromal cells but minimal effect on osteosarcoma cell growth. However, when delivered along with chemotherapeutics this stimulatory effect was completely abolished[7]. Thus, we concluded that BMP-2 delivery may not be an effective treatment option to induce bone formation within the damaged bone post-tumour resection while the patient is undergoing chemotherapy.

It is clear that chemotherapy can significantly hinder bone’s ability to regenerate[6c, 6d], yet all of the current biomaterial based studies investigating osteosarcoma and bone regeneration fail to validate how their therapy will fair while the patient is undergoing chemotherapy[9]. Furthermore, none of these studies fully interrogated the therapeutic potential of their implant at suppressing tumour growth whilst simultaneously aiding in bone regeneration[9], as two different models were utilized to evaluate their implant, either *in vitro* tumour cell eradication studies or a subcutaneous osteosarcoma tumour model (rather than an orthotopic model), and a separate segmental bone defect model in a healthy animal despite the fact that osteosarcoma has been shown to manipulate the physiological bone remodelling process[10]. Therefore, to truly understand the bone regenerative potential of an implant it is imperative to not only test its capabilities while the patient is undergoing chemotherapy but also to test its therapeutic potential in a disease state.

The controlled *in vivo* delivery of microRNAs (miRNAs), non-coding small RNAs that regulate gene expression, offer numerous therapeutic advantages and may potentially be developed to target both bone cancers and promote bone remodelling. miRNAs have been identified as gene expression master regulators and constitute an attractive target for treating cancer as they have been shown to regulate biological systems such as stemness, immunity and have been shown to play a crucial role in the initiation and progression of numerous cancers[11]. miR-29 consists of three mature members, miR-29a, miR-29b and miR-29c, which are encoded in two genetic clusters[12]. Members of this family, specifically miR-29b, has been shown to be silenced or down-regulated in many different types of cancer[13]. Subsequently, restoration of miR-29b has been found to elicit tumour-suppressive properties in a variety of cancers[13b, 13d, 13e, 14], but has not been studied in the context of osteosarcoma. It has also been shown that miR-29b plays a key role in bone remodelling by promoting osteoblast differentiation by downregulating TGF-β3 signalling[15], providing new insight into the use of miRNAs to induce osteogenesis. Furthermore, as miR-29b has also been shown to inhibit angiogenesis by targeting VEGF signalling pathways[16], it has the potential to aid in bone regeneration without enhancing vascularisation and thus tumour growth or metastasis. However, the therapeutic potential of miR-29b in osteosarcoma remains unknown. *In vitro* studies have shown that miR-29b delivery supresses proliferation and migration and induces apoptosis of osteosarcoma cells[17] and in some cases can sensitize the cells to chemotherapy[17b, 18]. Although informative, these studies used 2D *in vitro* cultures.

There are also several challenges that limit miRNA delivery as potential treatment option for osteosarcoma. These challenges include poor penetration of miRNAs into the tumour tissues, fast degradation time of unmodified miRNAs, and activation of the innate immune system leading to unexpected toxicities and undesirable side effects[19]. Injectable hydrogels as the miRNA delivery systems have become a research hotspot as it can efficiently avoid these problems by releasing the miRNA locally at the tumour site. By locally delivering the miRNA to the primary tumour site one can achieve superior transfection due to higher bioavailability, and reduced toxicity caused by non-specific uptake of the miRNA by normal healthy organs[19]. Other advantages of injectable hydrogels include their tuneable properties, controllable degradation, high water content, and shear thinning capabilities allowing them to be injected through a syringe to deliver miRNAs in a minimally invasive manner[20]. As hyaluronic acid (HA) is a biocompatible, non-immunogenic, and FDA-approved [21] and we have previously developed HA-based microneedle platform for non-invasive immunoregulation in skin transplants [22], we chose HA as our base hydrogel for the injectable delivery system. We also developed poly-beta-amino-esters (pBAE) nanoparticles as the intracellular delivery vehicle to deliver the miRNA as they have low toxicity and high biocompatibility due to their backbone of repeating ester groups that are biodegradable through hydrolysis in the cell cytoplasm[23]. Unlike lipid-based nanoparticles, which encapsulate the miRNAs within the liposome or micelle structure, pBAE polymeric chains electrostatically interact with the RNA therapeutics as the nanoparticle is being self-assembled, allowing for higher encapsulation and loading efficiencies[24]. Previous studies have demonstrated effective delivery of miR-29b via intertumoral injection or local administration of miR-29b via nanoparticle-mediated transfection[13b, 25]. Both studies demonstrated the therapeutic potential of localised miR-29b delivery at suppressing tumour growth in both prostate and cervical cancer[13b, 25]. Despite these promising results, no study has investigated the therapeutic potential of localised miR-29b delivery at suppressing tumour growth in osteosarcoma.

With this in mind, the overall goal of this study was to investigate if localised delivery of pBAE nanoparticles containing miR-29b to the primary tumour site, along with systemic delivery of chemotherapy, would suppress tumour growth whilst simultaneously providing the surrounding damaged bone the cues needed to normalise bone homeostasis even in the presence of chemotherapy (**Figure 1**). A pBAE nanoparticle delivery vector was developed and tested to efficiently deliver miR-29b to both osteosarcoma cells and surrounding stromal cells *in vitro* and *in vivo*. Using our previously developed *in vitro* spheroid model for osteosarcoma[7], we validated the therapeutic capabilities of miR-29b delivery in a controlled predictive model of the disease. Upon developing a pre-clinical metastatic murine model for osteosarcoma, we investigated the antitumour efficacy of localised delivery of miR-29b-complexes using a HA-based injectable system to enable efficient, local and sustained release of miR-29b to the primary tumour site. This allowed us to test the true therapeutic potential of the treatment in a diseased orthotopic model. The therapeutic potential was directly compared to and added as a potential add-on to the current clinical gold standard of systemic chemotherapeutic, Doxorubicin, to validate if the therapy can normalise the dysregulation of bone homeostasis caused by both the chemotherapeutics and osteosarcoma. Finally, as BMP-2 is known to be a strong inducer of bone remodelling[26], we directly compared our miR-29b treatment to BMP-2 delivery and validated its ability to normalise bone homeostasis and induce bone remodelling. We found that when miR-29b was delivered along with systemic chemotherapy, compared to chemotherapy alone, our therapy provided a significant decrease in tumour burden (45% reduction in tumour volume), increase in mouse survival (50% survival went from 24 days to 32 days), and a significant decrease in osteolysis (75% reduction in osteolysis) thereby normalising the dysregulation of bone lysis activity caused by the tumour. When directly compared to BMP-2 delivery, miR-29b significantly reduced osteolysis cause by the tumour and led to enhanced bone tissue distribution, even in the presence of chemotherapy. Together, our study highlights the therapeutic potential of miRNAs, specifically miR-29b, as a novel therapeutic add-on to conventional chemotherapy for targeting not only the primary tumour, but also normalise the dysregulation of bone homeostasis in the surrounding damaged bone.

**Figure 1:**
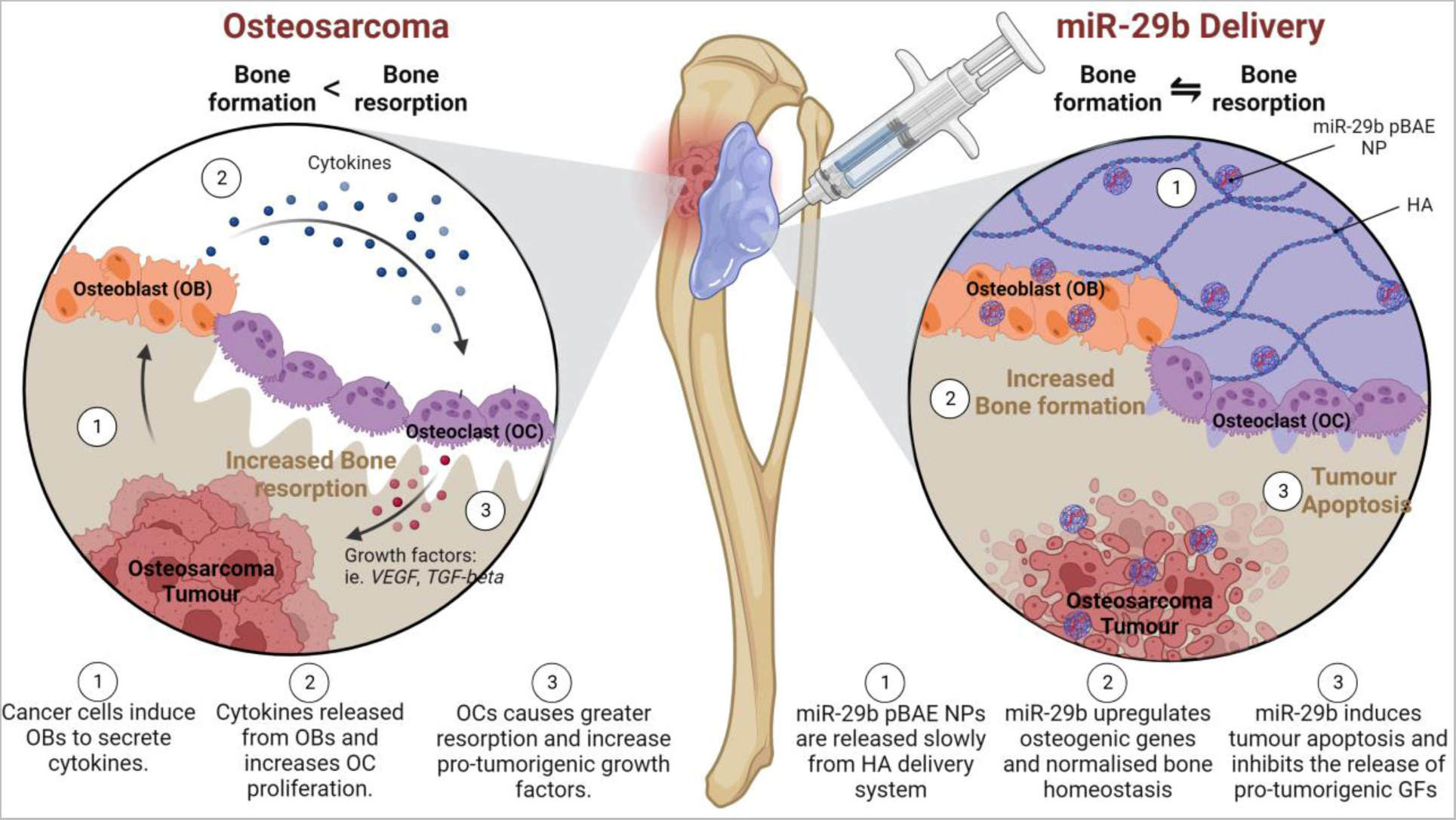
Schematic of the proposed dual-therapeutic role localized delivery of miR-29b:pBAE nanoparticles has in treating osteosarcoma. This will be achieved by inducing apoptosis in the tumour locally by restoring miR-29b expression within the tumour cells whilst simultaneously providing the surrounding damaged bone the cues needed to normalise the dysregulation of bone homeostasis caused by both the chemotherapeutics and the tumour (created with Biorender).

## Results

### Synthesis and characterization of pBAE nanoparticles

Synthesis of C6-CR3 polymer was previously reported[27]. C6-CR3 polymer was generated by reacting terminal acrylate groups from C6 polymer with arginine peptide (**Supplementary Figure 1 (A)**). C6-CR3 polymer was purified by precipitation and its molecular structure was characterized by ^1^H-NMR. The chemical structure of the resultant polymer was confirmed by the presence of signals associated with the conjugated arginine peptide (**Supplementary Figure 1 (B)**). We have previously confirmed that the C6-CR3 polymer was able to efficiently complex different types of nucleic acids, from small RNAi[23, 28] to DNA plasmids[29]. Here, the miR-29b complexation efficacy using C6-CR3 polymer was evaluated by agarose gel electrophoresis at different C6-CR3:miR-29b ratios (w/w). Complexation of C6-CR3 polymer with miR-29b revealed free miR-29b at ratios below 50:1 C6-CR3:miR-29b (w/w), while at a ratio of 50:1 or higher complete miR-29b complexation was observed (**Supplementary Figure 1 (D)**). At 100:1 ratio, the miR-29b nanoparticles were stable for more than 5 days in PBS with an average nanoparticle size of 151.2±1.8 nm. In contrast, at lower polymer:miR-29b ratios the nanoparticle stability was reduced (**Supplementary Figure 1 (E)**). Therefore, the 100:1 C6-CR3:miR-29b ratio was selected for the *in vitro* and *in vivo* experiments.

### PBAE nanoparticle-mediated transfection of human MSCs induces a pro-osteogenic and anti-angiogenic response whilst transfection of osteosarcoma cells (murine and human) induces a pro-apoptotic response alone

To analyse what effect miR-29b delivery would have on human MSCs, fluorescently labelled C6-CR3 polymer was complexed with either miR-29b or scramble miRNA (20nM, 8.3µg/mL) (see **Figure 2 (A)**). Following a 4-hour transfection with pBAE nanoparticles (Scramble; miR-29b), MSCs were cultured in osteogenic medium for 7 days, and then analysed for DNA content (*biocompatibility*), ALP Activity (*pro-osteogenic affect*), and VEGF release (*anti-angiogenic affect*). Confocal imaging validated cellular uptake of the pBAE nanoparticles 4 hours after transfection (see **Figure 2 (B)**). There was no significant difference in DNA content following miR-29b transfection, whereas there was a significant increase in ALP expression and a significant decrease in VEGF release when compared to the non-transfected and scrambled controls (see **Figure 2 (C)**).

**Figure 2:**
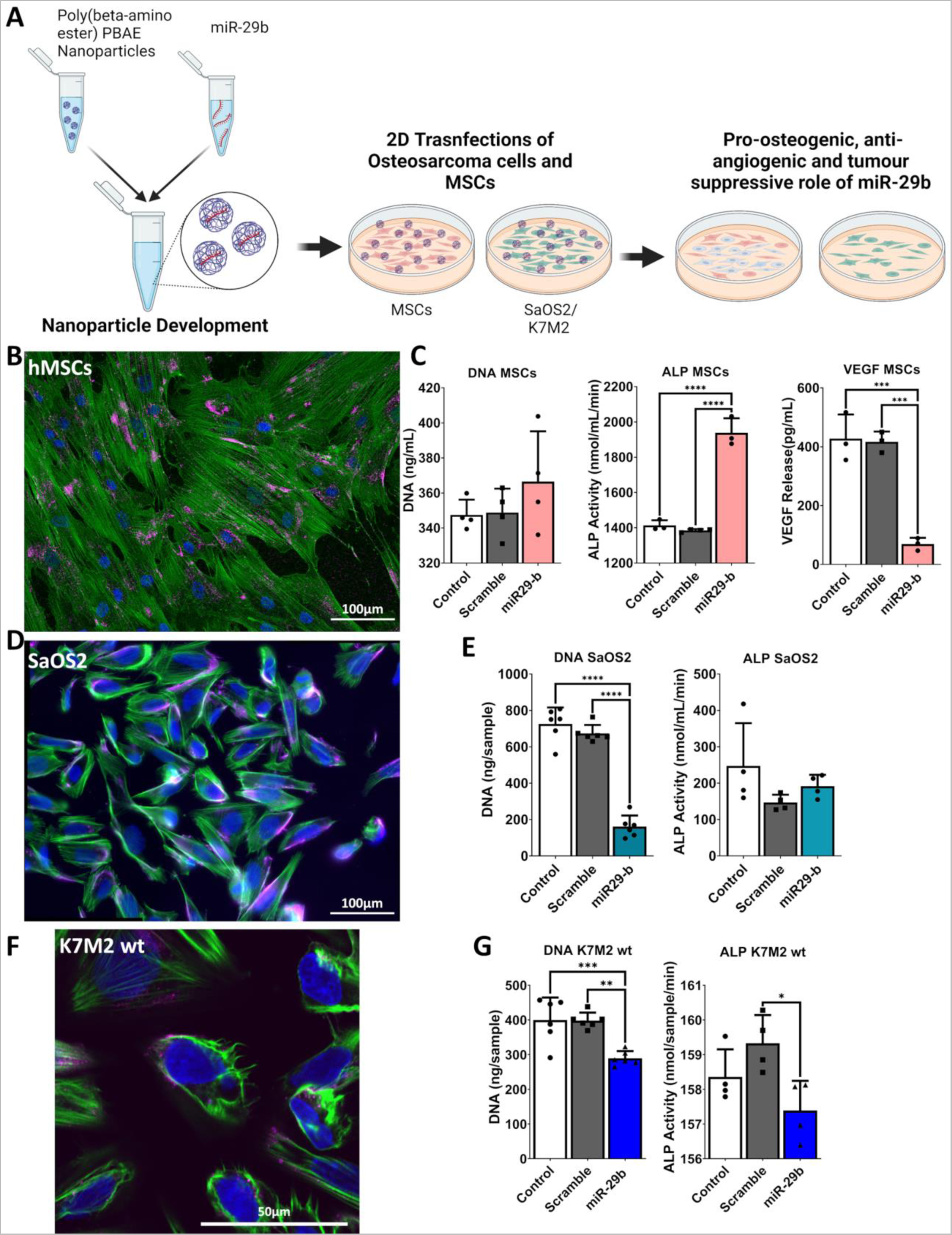
PBAE nanoparticle-mediated transfection of human MSCs and osteosarcoma cells in 2D culture. **(A)** Schematic of the experimental setup. Created with Biorender. **(B)** Confocal images of human MSCs following 4-hour transfection. **(C)** DNA Content, ALP activity and VEGF release over 7 days following transfection with miR-29b, or scramble loaded pBAE nanoparticles. **(D)** Confocal images of human osteosarcoma cells (SaOS2) following 4-hour transfection with fluorescently labelled-pBAE nanoparticles. **(E)** DNA Content and ALP activity over 3 days following transfection with miR-29b, or scrambled miR loaded pBAE nanoparticles. **(F)** Confocal images of Wild Type K7M2 mouse osteosarcoma cells following 4-hour transfection. **(G)** DNA Content and ALP activity over 3 days following transfection with miR-29b, or scramble loaded PBAE nanoparticles. In all confocal images nuclei are stained in blue, actin-cytoskeleton in green, and miR-29b pBAE nanoparticles in pink. All data is represented as Means ± SD *n*=4/6. Statistical differences were assessed using one-way ANOVA. *p<0.05, **>0.01, ***p < 0.001, ****p < 0.0001.

To understand what effect miR-29b delivery would have on osteosarcoma cells, human osteosarcoma cells (SaOS2) were transfected with pBAE nanoparticles loaded with either Scrambled miR or miR-29b. Following a 4-hour transfection period, the cells were cultured in osteogenic medium for 3 days, and then analysed for DNA content (*pro-apoptotic affect*) and ALP Activity. Confocal imaging validated cellular uptake of the pBAE nanoparticles 4 hours after transfection (see **Figure 2 (D)**). There was a significant decrease in DNA content following miR-29b transfection, when compared to the non-transfected and scrambled controls and no significant difference in ALP activity was observed between all three groups (see **Figure 2 (E)**). Osteosarcoma cells released negligible VEGF over the 3 days (data not shown).

Finally, a K7M2 cell line was chosen as it is considered highly aggressive with a reported pulmonary metastatic rate of over 90% in mice[30]. To ensure that miR-29b has a similar therapeutic response in murine cells as it does in human osteosarcoma cells, K7M2 cells were transfected with pBAE nanoparticles (Scramble; miR-29b). Following a 4-hour transfection period, K7M2 cells were further cultured in osteogenic medium for 3 days and analysed for DNA content and ALP Activity. Confocal imaging validated cellular uptake of the pBAE nanoparticles 24 hours after transfection (see **Figure 2 (F)**). There was a significant decrease in DNA content following miR-29b transfection, when compared to the non-transfected and scramble controls as previously seen with the human osteosarcoma cells (see **Figure 2 (G)**). There was also no significant difference in ALP activity between the control and the miR-29b or Scramble groups, however, there was a significant decrease in ALP activity between the Scramble and the miR-29b groups.

### Screening the therapeutic potential of miR-29b in a controlled predictive model of the disease

Recently, there has been increased interest in the use of 3D cultures to study the tumour microenvironment as they are more predictive of the *in vivo* situation[7, 31]. With this in mind, osteosarcoma tumour spheroids containing a co-culture of both MSCs and osteosarcoma (SaOS2) cells were generated using a hydrogel microwell system[7] (see **Figure 3 (A)**). Following a 4-hour transfection with pBAE nanoparticles (miR-29b, 20nM), tumour spheroids were further cultured in osteogenic medium ± the FDA approved chemotherapeutic Doxorubicin (1.8µM, the IC_50_ concentration for SaOS2 cells was as previously determined[7]) for 7 days. Tumour spheroid growth was significantly hindered as evidenced by a significant decrease in DNA content regardless of transfection. In the absence of chemotherapeutics, miR-29b transfection led to a significant increase in DNA content (see **Figure 3 (B)**). Flow cytometry data revealed that this increase in DNA content was due to an increase in MSC proliferation or selective osteosarcoma cell apoptosis as the ratio of MSCs:SaOS2 cells tripled following transfection (see **Figure 3 (C)**). This was further verified in the H&E staining with significant increase in ECM production, tumour spheroid size, and nucleic staining present in the miR-29b group (see **Figure 3 (D)**). The cytotoxic effect of the Doxorubicin on tumour progression was not hindered with the addition of miR-29b transfection, as evidenced by reduced tumour spheroid size and minimal nucleic staining present in the groups cultured in Doxorubicin.

**Figure 3:**
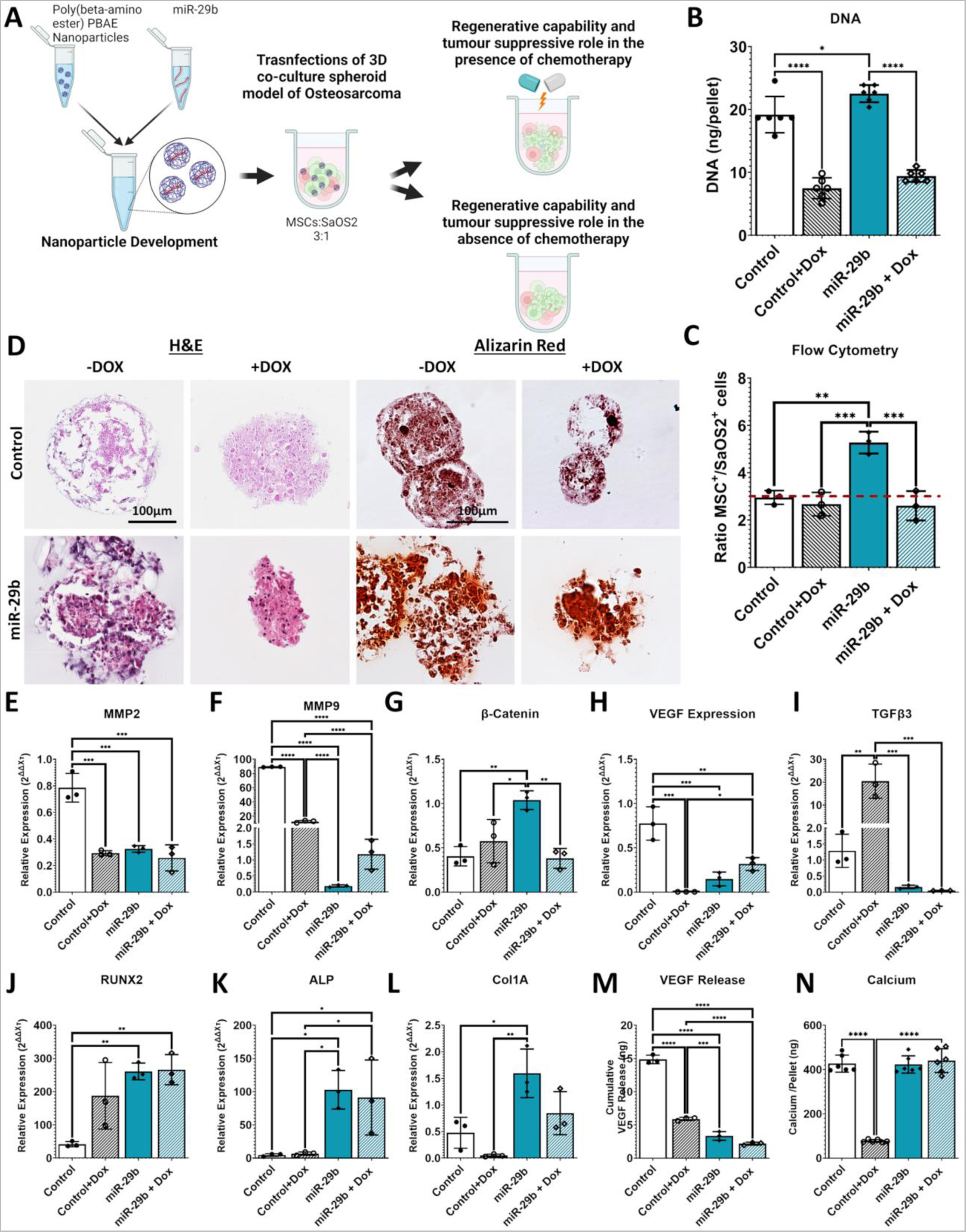
Spheroid Growth, Cancer Progression, and Mineralization following miR-29b transfection in the presence/absence of Doxorubicin using 3D tumour spheroid model of osteosarcoma. **(A)** Schematic of the experimental setup. Created with Biorender. **(B)** DNA content and **(C)** analysis of the ratio of MSCs+ to SaOS2+ cells present within the spheroids. **(D)** H&E and Alizarin Red staining of the tumour spheroids following 7 days post-transfection. Images taken at 20X. **(E-L)** mRNA levels of *MMP2, MMP9, β-Catenin, VEGF, TGFβ3, RUNX2, ALP* and *Col1A* were analysed by qRT-PCR. **(M)** VEGF Release and **(N)** Calcium Content present within the tumour spheroids. All data was obtained following a 7-day culture period post-transfection and is represented as means ± SD *n* = 6 (25 spheroids per experimental replicate) for biochemical assays, *n* = 3 (100 spheroids per experimental replicate) for PCR analysis. Statistical differences were assessed using one-way ANOVA with Tukey post-test. *p < 0.04, **p < 0.007, ***p < 0.0005, ****p < 0.0001.

To understand the mechanism by which miR-29b delivery induces selective apoptosis in the cancer cells, as seen in our 2D culture and flow cytometry data, we analysed four well-characterised apoptotic genes *BCL-2*, *BAX*, *CASP 9* and *CASP 3* (see **Supplementary Figure 2**). MiR-29b delivery led to a significant decrease in *BCL-2* expression in addition to a significant increase in *CASP 9* expression. This demonstrates that one potential mechanism in which miR-29b delivery induces apoptosis is by initiating the intrinsic apoptotic pathway.

Finally, we investigated what effect miR-29b transfection was having at a gene level within our novel spheroid model. Looking at the prognostic genes for osteosarcoma, there is a clear antitumour effect when miR-29b was delivered to the tumour model. This effect was equivalent to the addition of Doxorubicin, as evident by the significant reduction in known prognostic markers *MMP2* and *MMP9*[32] mRNA expression in *miR-29b* group compared to the *control* (see **Figure 3 (E, F)**). Interestingly, there was a significant increase in *β-Catenin* mRNA expression when miR-29b was delivered alone (see **Figure 3 (G)**). As *β-Catenin* signalling also plays an important role in osteogenesis this increase in expression may be due to the increased osteogenic differentiation of the MSCs. MiR-29b delivery was seen to have an anti-angiogenic effect regardless of the addition of Doxorubicin. This was seen at both the gene (*VEGF expression*) and protein level (*VEGF release*) (see **Figure 3 (H, M)**). There was a significant decrease in *TGF-β3* mRNA expression regardless of the addition of Doxorubicin when the spheroids were transfected with miR-29b. As studies have shown that one of the mechanisms behind miR-29b ability to induce osteoblast differentiation is by downregulating TGF-β3[15], it validates the transfection efficiency of the pBAE nanoparticles to deliver the miR-29b even in a 3D environment (see **Figure 3 (I)**). Finally, as previously seen in our 2D *in vitro* culture studies, alizarin red staining revealed that miR-29b transfection led to a significant increase in mineralisation in our tumour spheroids. This was apparent even when the tumour spheroids were also cultured in the presence of Doxorubicin (see **Figure 3 (D)**). This was further verified at both the gene level (*RUNX2, ALP, Col1A* mRNA expression) and protein level (calcium content) (see **Figure 3 (J-L, N)**), verifying miR-29b delivery promotes bone remodelling.

### Development of hyaluronan-based injectable system for the local and sustained release of pBAE:miR-29b nanoparticles

After demonstrating the dual therapeutic role miR-29b delivery has *in vitro*, we tested its efficacy at suppressing tumour growth whilst simultaneously normalising the dysregulation of bone homeostasis caused by both the chemotherapeutics and the osteosarcoma within the surrounding bone. A novel hyaluronan (HA)-based injectable system was developed to enable efficient, local and sustained release of either the pBAE:miR-29b nanoparticles or BMP-2 (see **Figure 4 (A)**).

**Figure 4:**
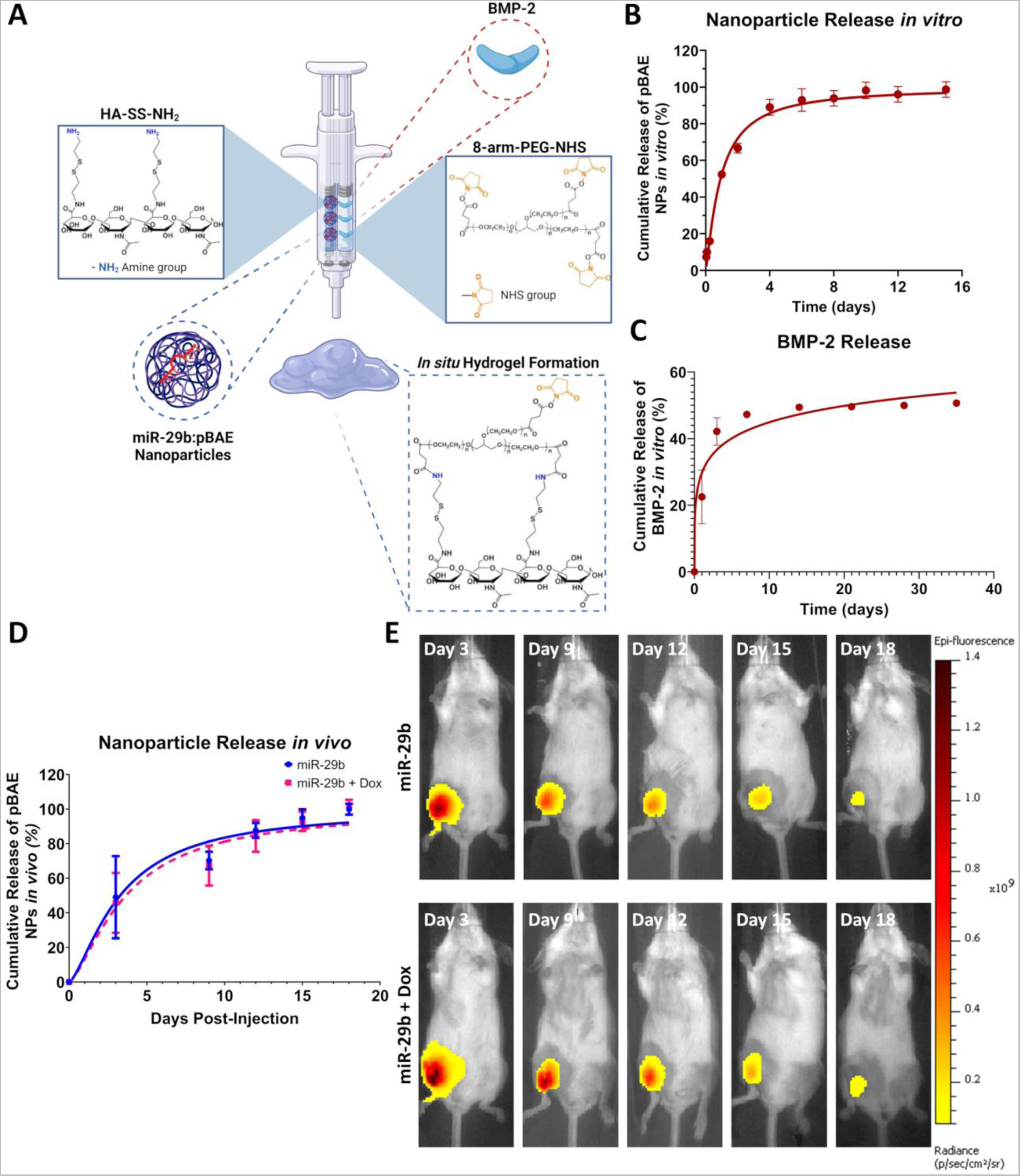
Development of hyaluronan-based injectable system for the local and sustained release of pBAE nanoparticles or growth factors. **(A)** HA-injectable hydrogel fabrication process: aqueous amine-modified HA (HA-SS-NH2) solution is crosslinked using the NHS-terminated 8-arm PEG crosslinker. The final chemical structure of the injectable HA hydrogel is found in the final scheme. Created with Biorender. **(B)** Cumulative release of pBAE nanoparticles from HA hydrogel over 15 days *in vitro*. **(C)** Cumulative release of BMP-2 from HA hydrogel over 35 days *in vitro*. **(D, E)** Cumulative release of pBAE nanoparticles from HA hydrogel over 18 days following local administration of the HA hydrogel in orthotopic model for osteosarcoma. All data is represented as Means ± SD *n*=4 for *in vitro* release and *n*=8 for *in vivo* release.

To form the HA-based hydrogel, HA primary amines were reacted with the 8-arm-PEG-NHS crosslinker containing a succinimidyl functional group, allowing for spontaneous hydrogel formation. The crosslinking speed of the hydrogel approximated around 1-2 minutes, thereby allowing for the delivery system to be easily injected using a standard 25G needle. *In vitro* release kinetics revealed that it took about 10 days for 100% of the nanoparticles to be released from the hydrogel (see **Figure 4 (B)**). As HA-based hydrogels have been successfully tested as injectable carriers of bone morphogenetic protein-2 (BMP-2)[33], which is widely known to induce bone remodelling[26] it was chosen as the control to evaluate the pro-osteogenic potential of our miR-29b:nanoparticles. *In vitro* release kinetics of the BMP-2 from the same delivery vehicle revealed a burst release of BMP-2 in the first three days with 42% of the overall concentration of growth factor released in this time frame (see **Figure 4 (C)**), over the next 33 days in culture there is a slow release of BMP-2 with only another 10-12% release of the total growth factor during this time.

Next, using our newly developed metastatic murine model for osteosarcoma (see **Supplementary Figure 5**), we validated the injectability our HA delivery system to non-invasively deliver the pBAE nanoparticles to the primary tumour site. The HA-based injectable system crosslinked *in situ* and allowed for local and sustained delivery of the miR-29b to the primary tumour site. Florescent IVIS imaging verified the *in vitro* release kinetics profile with ∼100% of the nanoparticles released from the HA hydrogel in ∼15-18 days post-implantation (see **Figure 4 (D)**). IVIS imaging also demonstrated that when locally administered there was focused delivery of miR-29b to the primary tumour site as evident by the accumulation of the nanoparticles around the tumour, with no downstream signal for any healthy organs (see **Figure 4 (E)**).

### Localised miR-29b delivery supressed tumour growth locally and when combined with systemic Doxorubicin significantly increased survivability

Next, we assessed whether a combination therapy (systemic doxorubicin and localised miR-29b) would supress tumour growth compared to either therapy alone using an orthotopic model for osteosarcoma (see **Figure 5(A)**). Systemic Doxorubicin of 2 mg/kg was administered intravenously twice a week over the course of 3 weeks and is equivalent to a dose of 6.5 mg/m^2^ given clinically[34]. Tumour size was measured by luminescence and vernier callipers twice a week for three weeks (see **Figure 5 (B, C)**). MiR-29b-treated mice (*miR-29b, miR-29b + Dox*) showed significantly slower tumour growth than all three controls (see **Figure 5 (B, C)**). When miR-29b was delivered along with systemic chemotherapy, compared to chemotherapy alone, our therapy provided a 45% reduction in tumour volume over 15 days. This corresponded to a significant increase in survival when compared to all three controls (see **Figure 5 (D)**). Specifically when miR-29b was added to conventional chemotherapy there was an increase in the 50% survival - from 24 days (chemotherapy alone) to 32 days (combination therapy). Furthermore, it was directly due to the miR-29b delivery and not the HA hydrogel, as the hydrogel alone group had no effect on tumour growth or survivability (see **Figure 5 (D)**).

**Figure 5:**
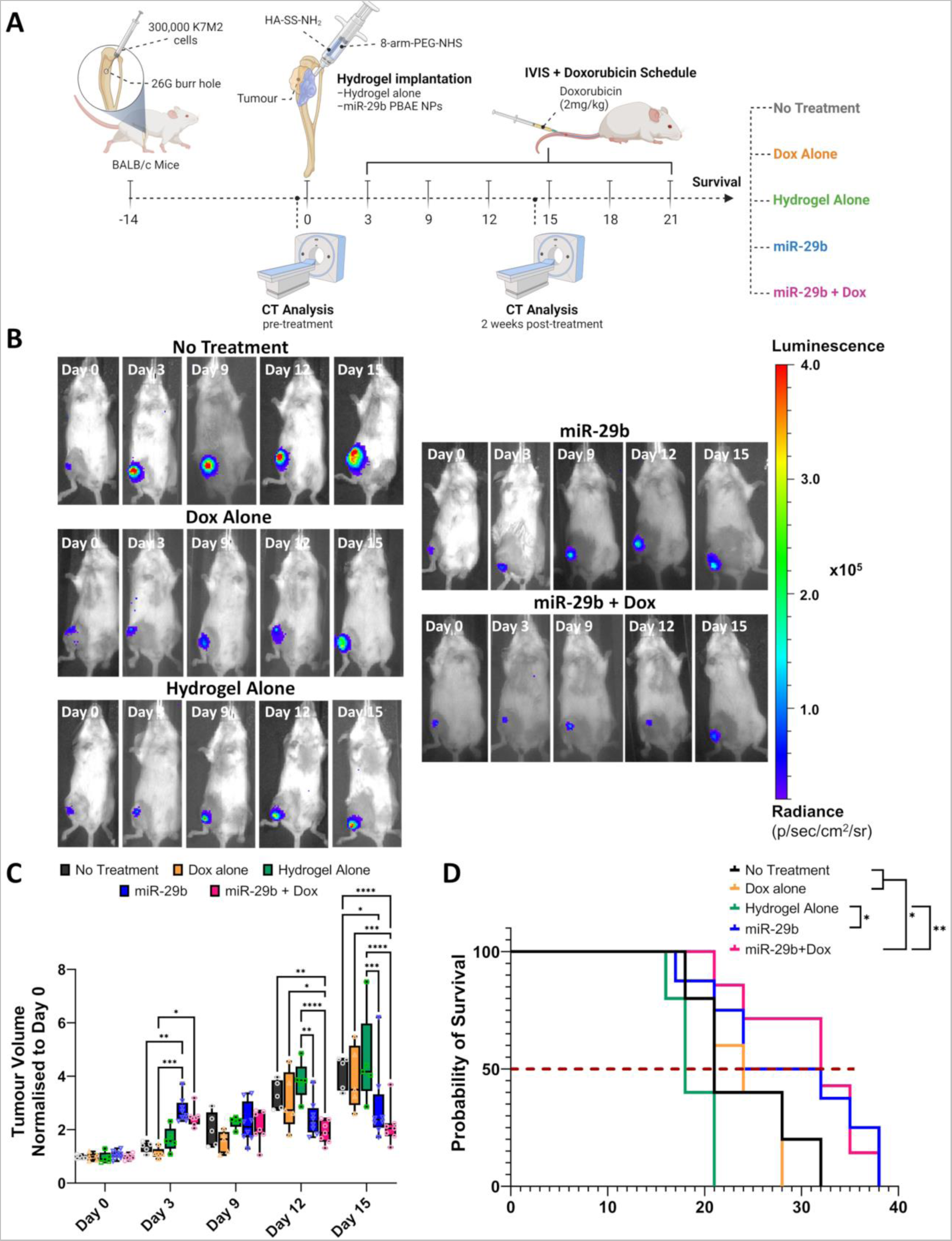
Localised miR-29b delivery supressed tumour growth locally and when combined with systemic Doxorubicin significantly increased survivability. **(A)** Schematic of the developed metastatic orthotopic osteosarcoma mouse model including all the treatment groups. Created with Biorender **(B)** Live imaging (IVIS) following inoculation with K7M2-Luc labelled osteosarcoma cells. **(C)** Quantitative evaluation of tumour size calculated using the following formulae (Width^2^xLength)/2 at different time points. All data was obtained over 15 days following the various treatment regimens and is represented as means ± SD (*n*=5/8 per group). Statistical differences were assessed using two-way ANOVA with Tukey post-test. *p < 0.01, **p < 0.006, ***p < 0.0008, ****p < 0.0001 **(D)** Kaplan–Meier survival curves of mice treated with the indicated formulation using a 1000mm^3^ tumour volume or poor body condition as the endpoint criterion. Statistical analysis was performed using a log-rank Mantel–Cox test. *p < 0.02, **p < 0.0018. Red dashed line denotes 50% survival rate.

### Localised delivery of miR-29b significantly reduced osteolysis caused by chemotherapeutics and the primary tumour and normalised bone homeostasis within the surrounding damaged bone following two weeks of treatment

We sought to assess what effect if any the local delivery of miR-29b would have on the surrounding damaged bone. Our *in vitro* studies demonstrated the miR-29b delivery promoted bone remodelling by inhibiting TGF-β3 signalling thereby significantly increasing the expression for a panel of well-established osteogenic genes. Our *in vivo* study cooperated with these findings as µCT imaging, specifically around the primary tumour site (see **Figure 6 (B)**, red box), revealed a significant increase in bone volume when compared to systemic chemotherapeutic alone or untreated controls. Thereby, verifying that miR-29b delivery normalises the bone homeostasis within the surrounding damaged bone by normalising the dysregulation of bone lysis activity caused by the growing tumour. A similar trend was seen but without significance, in the same group when combined with systemic chemotherapeutics (p = 0.066). There was on average 75% reduction in osteolysis when miR-29b was delivered along with systemic chemotherapy, compared to chemotherapy alone. MicroCT 3D reconstructions and histological analyses revealed the extent of the osteolysis due to growing tumour in the untreated controls (*No treatment, Hydrogel Alone*) and clinical standard (*Dox Alone*), see **Figure 6 (C, D)**. In the untreated controls and clinical gold standard it clear that a sizable part of the tibia has been lysed away in place of tumour, due to the dysregulation of bone homeostasis caused by the tumour. Furthermore, there is little to no marrow cavity remaining in these groups (see **Figure 6 (D)**; denoted by M). On the other hand, both miR-29b treated groups (*miR-29b* and *miR-29b + Dox*) show slightly reduced tumour burden with enhanced bone tissue distribution complete with functional marrow cavities.

**Figure 6:**
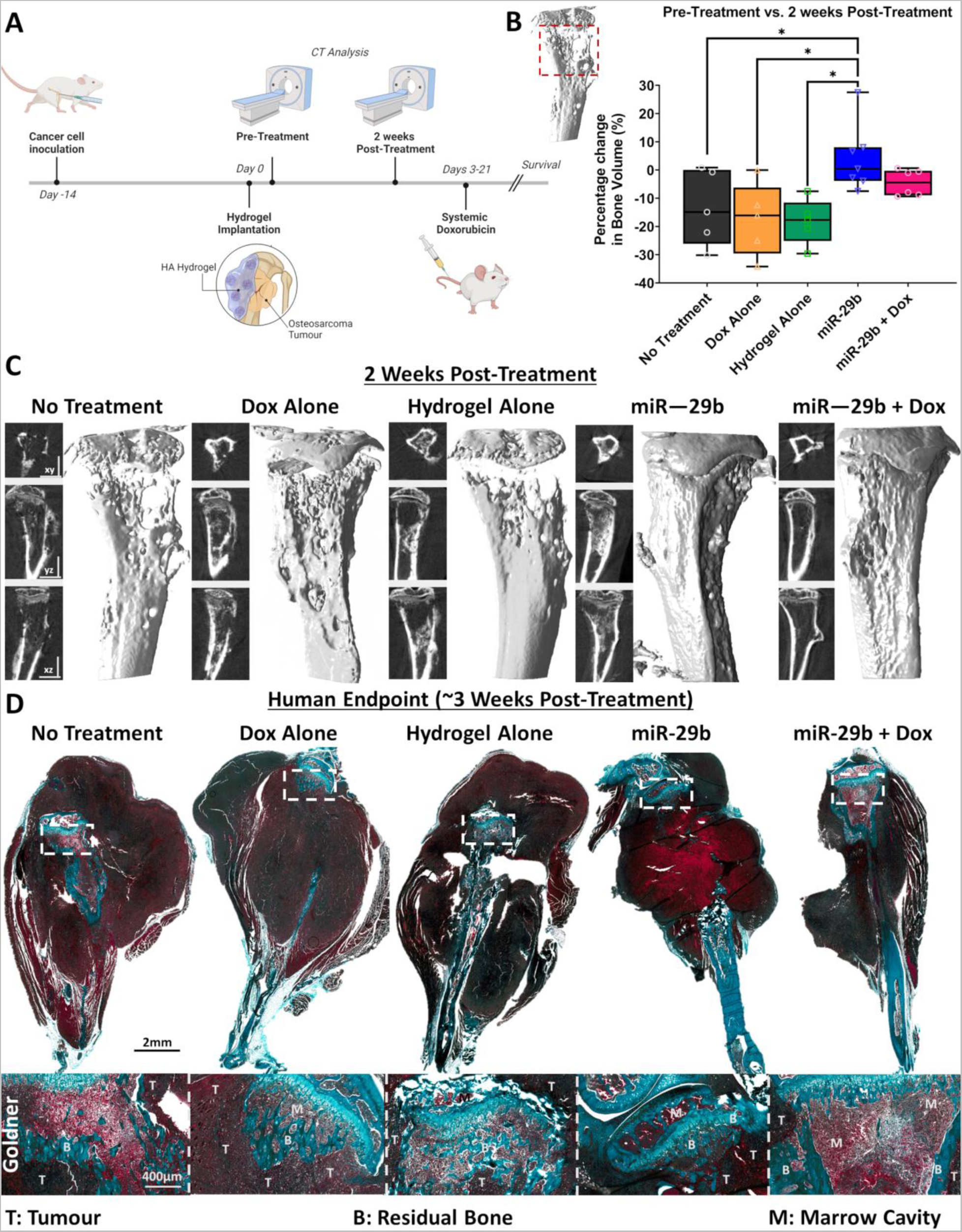
MiR-29b delivery significantly reduced osteolysis due to tumour burden and led to an increase in bone volume after 2-weeks of treatment. **(A)** Schematic of the timeline for µCT analysis to assess osteolysis following treatment with miR-29b:nanoparticles. **(B)** Percentage change in bone volume pre vs. post treatment. All data is represented as Means ± SD *n*=5/8. Statistical differences were assessed using a one-way ANOVA with Tukey post-test. *p < 0.01. **(C)** 3D reconstructions and x-rays images in the xy,yz,zx planes of the worst healers of the µCT data for each group at following 2 weeks of treatment. **(D)** Representative Goldner’s Trichrome– stained sections from all the experimental groups which mice all had a similar endpoint (∼3 weeks post-treatment). Images were taken at 20X. B denotes remaining bone tissue, M denotes marrow and T denotes tumour.

### Systemic doxorubicin and localised miR-29b delivery significantly reduced osteolysis and better maintained normal bone homeostasis in the long-term compared to local BMP-2 delivery

To validate the clinical relevance of localised delivery of miR-29b to induce bone remodelling and normalise bone homeostasis, the therapeutic potential was directly compared to BMP-2 delivery, which is widely known to induce bone remodelling[26]. A concentration of 5µg of BMP-2[35], was loaded into the in the HA-based injectable delivery system and following the 14-day inoculation period was non-invasively injected around the primary tumour site. MicroCT analysis was used to visualize and quantify bone formation around the primary tumour site pre-treatment (which corresponds to 2-weeks after tumour induction), 2- and 4-weeks post-treatment (which corresponds to 4- and 6-weeks after tumour induction respectively). There was a trend towards increase bone formation in both the *BMP-2* and *miR-29b* groups (p = 0.5) compared to their counterparts with combined systemic chemotherapeutics (*BMP-2 + Dox, miR-29b + Dox*) 2-weeks post-treatment (see **Figure 7 (B)**). MicroCT 3D reconstructions reveal that this increase in bone volume is due to the presence of ectopic bone formation surrounding the tibia. This was not present in the *miR-29b + Dox* group (see **Figure 7 (D)**). Interestingly, there was more ectopic bone formation observed surrounding the tibia (denoted by red arrows) in the *BMP-2* group when compared to the *BMP-2 + Dox* group even though both groups received the same hydrogel with the same concentration of BMP-2, the only difference was the addition of systemic Doxorubicin.

**Figure 7:**
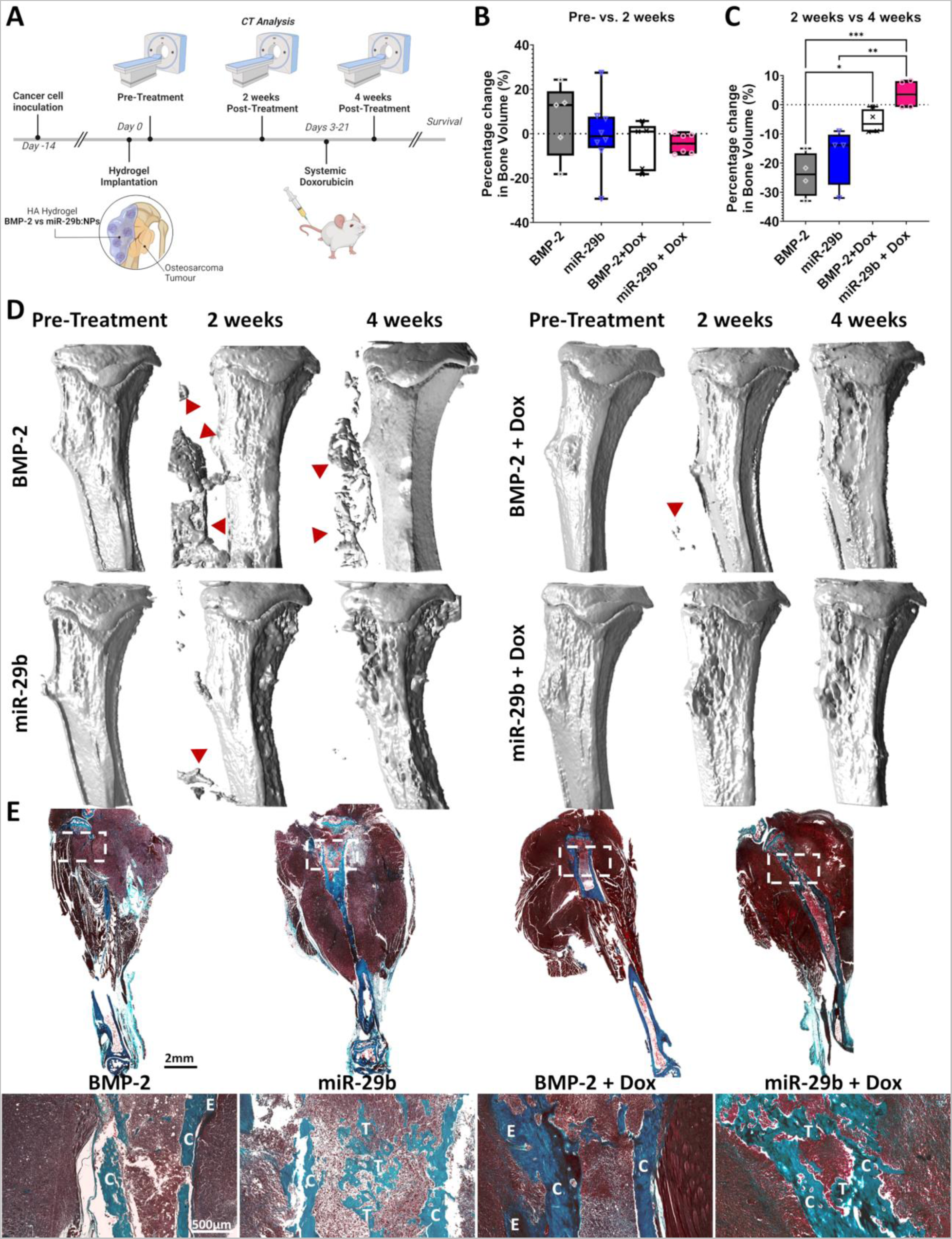
Localised delivery of miR-29b significantly reduced osteolysis and led to enhanced bone tissue distribution compared to clinical gold standard BMP-2 delivery even in the presence of chemotherapeutics. **(A)** Schematic of the timeline for µCT analysis to assess osteolysis following treatment with miR-29b:nanoparticles vs BMP-2 delivery. Percentage change in bone volume **(B)** pre-vs. 2 weeks post-treatment and **(C)** 2 weeks vs. 4 weeks post-treatment for all four treatment groups. All data is represented as Means ± SD *n*=5/8 (pre and 2-week post treatment), *n*=4 (4-weeks post treatment). Statistical differences were assessed using a one-way ANOVA with Tukey post-test. *p < 0.01, **p<0.004, ***<0.0007. **(D)** 3D reconstructed *in vivo* µCT analysis of the mouse tibias over the 4 weeks of treatment. Red arrow heads denote ectopic bone. **(E)** Representative Goldner’s Trichrome–stained sections from all the experimental groups which mice all had a similar endpoint. Images were taken at 20X. C denotes remaining cortical bone, T denotes remaining trabecular bone, E denotes ectopic bone formation.

Despite the small number of animals which survived to the 4-week timepoint (*n* = 4 per group), a clear effect of the combination therapy on osteolysis was observed. There was an increase in osteolysis (measured by percentage change in BV/TV) in both the *BMP-2* and *miR-29b* groups from 2- to 4-weeks post-treatment (see **Figure 7 (C)**). In contrast, bone homeostasis was maintained in the *BMP-2 + Dox* and the *miR-29b + Dox* groups as bone volume remained the same in the *BMP-2 + Dox* group, while there was a significant increase in bone formation as bone volume was significantly increased from 2- to 4-weeks post-treatment in the *miR-29b + Dox* group. This was further verified in the 3D reconstructions and histological analyses (see **Figure 7 (D, E)**). A significant amount of ectopic bone formation was present even 4-weeks post-treatment in the *BMP-2* group. MicroCT reconstructions showed little change to the cortical bone in the BMP-2 group but histological analysis revealed little to no trabecular bone present within the cortical shell due to tumour growth. The overall bone distribution was better maintained in the *miR-29b* group compared to the *BMP-2* group, as both cortical and trabecular bone were still present. A similar trend was seen in the corresponding groups with combined systemic chemotherapeutics (*BMP-2 + Dox*, *miR-29b + Dox*). Looking at the bone architecture we can see a small amount of ectopic bone formation with only the cortical shell remaining in the *BMP-2 + Dox* group. Conversely in the *miR-29b + Dox* group superior bone distribution and architecture is observed in both the cortical and trabecular bone (see **Figure 7 (E)**).

## Discussion

Despite the distinct clinical need for more effective treatment options, there has been no major improvements in the treatment of osteosarcoma since the 1970s[3]. Furthermore, as osteosarcoma is such an aggressive disease, the surgical intervention usually involves total reconstructions of the limbs or in most cases amputation, despite this, most of the research to date has focused on the prevention of metastases, with little attention to bone repair or salvation. This study presents a novel gene therapy as a potential add-on to the current gold-standard treatment for osteosarcoma, to not only significantly reduce tumour burden but also aid in the repair of the surrounding damaged bone by normalising the dysregulation of bone lysis activity caused by the chemotherapeutics and tumour. Here, we developed a pBAE nanoparticle delivery vector and tested its ability to efficiently deliver miR-29b to human and murine osteosarcoma cancer cells and surrounding healthy stromal cells *in vitro* in both 2D and 3D culture. Using our *in vitro* co-culture spheroid model for osteosarcoma[7], we confirmed the dual therapeutic capabilities of miR-29b delivery in a controlled predictive model of the disease. Finally, in an orthotopic metastasis model for osteosarcoma, we demonstrated the antitumour efficacy of localised delivery of pBAE:miR-29b complexes using a novel HA-based injectable delivery system. As we hypothesised that miR-29b may have a dual therapeutic role, we also verified the bone regenerative capabilities of miR-29b delivery versus traditional BMP-2 delivery and confirmed its ability to normalise bone tissue homeostasis, and in fact promote bone formation, which BMP-2 delivery failed to do. Furthermore, we not only validated the therapeutic capabilities of our miR-29b:complexes to repair the damaged surrounding bone while the patient is undergoing chemotherapy but also tested its therapeutic potential in a disease state, which all other biomaterial based studies have failed to do[9]. Taken together, the results from this study demonstrate not only the therapeutic potential of localised miRNA delivery in osteosarcoma tumour suppression but also its potential to aid in bone repair post-tumour resection for all types of bone cancers.

MicroRNAs (miRNA) have emerged as potential therapeutic targets or tools for cancer treatment, yet there are still delivery limitations that impede the advancement of miRNA therapy. This is largely due to low delivery efficiency as a result of poor penetration of miRNAs into the tumour tissues. Herein, we developed a pBAE nanoparticle delivery vehicle that was significantly more effective at delivering miRNA than commercially available transfection reagent Lipofectamine (**Supplementary Figure 6**). This increase in transfection efficiency is due the cationic nature of the nanoparticles, which enables them to condense the negatively-charged miRNA more efficiently than other polymeric based nanoparticles[29, 36]. Additionally, the cationic surface density (5.6mV ± 3.5) of pBAE nanoparticles also allows for efficient cell internalization which improves their endosomal escape due to the proton sponge effect. Furthermore, the pBAE backbone polymer contain ternary amine groups which acts as a buffer in the low pH environment of endosomes, which further leads to its disruption, and release of the encapsulated miRNA, resulting in more efficient delivery than liposomal-mediated transfection[29, 36]. In addition, our pBAEs were modified with peptides moieties to further increase their biocompatibility and cell internalization[27, 29, 37].

Next, we tested the penetrability of the pBAE particles to deliver the miR-29b in a 3D spheroid model of osteosarcoma[7]. As we needed the *in vitro* model to mimic the *in vivo* tumour microenvironment as much as possible, we incorporated both the osteosarcoma cells and the surrounding stromal cells within the model[7]. The 3D spheroid model also allowed for 3D spatial arrangement of the two cell types with enhanced cell to cell contact and the ability to form proliferative gradients, hypoxia, and necrosis[7, 38]. Furthermore, certain ECM components were found to be expressed at high levels in 3D spheroids, hence they have the potential to establish the penetration barriers seen *in vivo*, thereby allowing us to study the penetration, distribution, and uptake of the nanoparticles within these models[38–39]. Previous studies have suggested that nanoparticles which are capable of diffusing through pores between the collagen fibrils, measured between 20–40 nm, are needed to penetrate compact tumours, or between 75–130 nm in size when penetrating poorly organized tumours[38, 40]. Yet, despite their larger size (151 nm), our pBAE nanoparticles were capable of penetrating our tumour spheroids (∼0.28mm in diameter[7]) after only 4 hours of transfection, resulting in measurable changes at the genomic level within the two cell types (**Figure 2**). This increase in penetration may also be attributed to the cationic nature of the pBAE nanoparticles. Previously, studies have shown that cationic nanoparticles are significantly more effective in tumour growth inhibition than their anionic or neutral counterparts in a variety of tumour models due to improved permeability and penetration capabilities[41]. Furthermore, as the current process for testing the effectiveness of nanoparticles relies greatly on animal models, our unique spheroid system provides useful insights into the effectiveness of the pBAE nanoparticles in delivering the miR-29b within the tumour microenvironment before conducting a full pre-clinical *in vivo*study.

Dysfunction of miRNAs is often associated with tumour formation and progression as manipulation of the oncogenic pathways can influence tumour progression[19]. Specifically, studies have shown that miR-29b was significantly down-regulated in osteosarcoma tissues[42]. Previous *in vitro* studies have shown that miR-29b delivery to osteosarcoma cells supresses proliferation and migration and induces cell apoptosis of osteosarcoma cells[17]; however, the mechanism by which miR-29b induces apoptosis in osteosarcoma cells remains unclear. Our current study establishes that the delivery of miR-29b to the osteosarcoma cells, where miR-29b is down-regulated or silenced, induces the intrinsic apoptotic pathway. Briefly, miR-29b downregulates the anti-apoptotic protein B-cell lymphoma (*Bcl-2*), resulting in mitochondrial dysfunction and subsequent release of cytochrome c and activation of caspases-9 (*CASP 9*) (see schematic in **Supplementary Fig. 2**). Interestingly, when miR-29b is delivered to healthy cells, where its expression is normal, it led to an upregulation of miR-29b, which had the opposite effect. When miR-29b was delivered to the surrounding stromal cells, it induced cell proliferation and promoted osteogenic differentiation as evident in the flow cytometry and PCR data (see **Figure 2**). There are many studies which have characterized the role miR-29b plays in relation to tumorigenesis[13], but only a few studies which have examined its role in the osteogenesis[15a, 43]. Specifically, studies have shown that miR-29b is a positive regulator of osteoblast differentiation by down-regulating inhibitory factors of osteogenic signalling pathways and controlling expression of collagen in differentiated osteoblasts[15a]. Our current study follows on from this by delineating the network of genes regulated by miR-29b and demonstrating how this stimulatory effect is unaffected by the addition of chemotherapeutics. Taken together, our study confirms the dual therapeutic role of miR-29b delivery by enabling selective cancer-cell killing while simultaneously promoting osteogenesis in the surrounding healthy tissues.

In 2019, under the organization of the Bone Tumour Biology Committee of the Children’s Oncology Group, a team of clinicians and scientists sought to define the state of the science and to identify questions that, if answered, have the greatest potential to drive fundamental clinical advances for osteosarcoma[44]. Here, they suggest that transformative advances in the treatment of osteosarcoma will not likely come from intensifying chemotherapy, but on developing truly novel therapeutic approaches[44]. Our study demonstrates that i*n vivo* delivery of miR-29b carried on pBAE nanoparticles which are embedded in a HA delivery system and injected adjacent to the tumour, allowed for local and sustained release of the miRNAs and efficiently slowed down tumour growth in an orthotopic mouse model, with no systemic side effects (see **Figure 4-5**). This therapeutic effect has previously only been shown in prostate and cervical cancer[13b, 25], never in osteosarcoma. Moreover, delivery of the combined therapy (systemic doxorubicin and localised miR-29b) further improved preclinical outcomes, leading to a significant increase in 50% survival from 24 days to 32 days when compared to the current clinical gold standard of systemic chemotherapeutics. This validates the plausibility of introducing our novel localised miRNA therapy as a potential add-on to conventional chemotherapy.

Despite the promising results in reducing the tumour burden locally, none of the animals were fully cured. This may in part be attributed to the clinically relevant orthotopic model used in this study. Over the last 5 years there are over 20 journal articles[9] investigating the divergent relationship between osteosarcoma elimination and bone regeneration, yet none of these studies evaluated the dual-therapeutic potential of their therapy in an orthotopic model. Instead they utilise two different *in vivo* models, a subcutaneous osteosarcoma tumour model (rather than an orthotopic model), and a separate segmental bone defect model in a healthy animal. Despite the fact and osteosarcoma has been shown to manipulate the physiological bone remodelling process[10] so analysing its ability to regenerate bone in a healthy animal is not a true representation of what would happen if applied in the clinic. Only one research group has come close by utilising a tumour plague embedment method[45]. Yet, this method still utilised a subcutaneous tumour model which does not take account of the proper interactions between the tumour cells and their normal bone/muscle/cartilaginous microenvironment. This study is the first to evaluate the dual-therapeutic effect in a clinically relevant orthotopic model for osteosarcoma.

Another mitigating factor may be the dosing regimen used in this study. Only one dose of miR-29b was administered and although there was a therapeutic response in the 15 days in which the nanoparticles were being released, this did not lead to full tumour elimination. Future studies aim to evaluate if a second dose of miR-29b after 15 days may significantly increase its therapeutic potential and prevent metastases. Furthermore, the interaction of tumour and host immune system has long been recognized to be a critical aspect of tumour survival in malignancies. Despite this, the relationship between tumour and host immune microenvironment, their interactions, and opportunities for exploitation of immune-mediated therapies remain poorly defined in osteosarcoma[44]. Previous studies have shown that chemotherapeutics like cisplatin and doxorubicin, have the capacity to upregulate programmed death-ligand1 (PD-L1) expression on cancer cells and which in turn promotes antitumor immunogenicity, via activation of cytotoxic T cells[46]. We have also previously shown in our lab that delivery of Stimulator of Interferon Genes (STING) agonist is effective therapy at eliminating melanoma[47]. As metastasis was not the focus of this study, we did not investigate any further combination therapies but future studies will evaluate the synergistic potential of a triple-therapy of combining localised nanoparticle-mediated gene delivery with systemic chemotherapy and immunotherapies like anti-PD1 or STING. Finally, although chemotherapy is effective in controlling cancer cell growth, it has also been shown clinically to disrupt bone homeostasis[6], resulting in a dysregulated bone lysis activity significantly hindering the surrounding bone’s ability to regenerate following surgical intervention. Previously, we suggested that BMP-2 delivery may not be an effective therapy to aid with the regenerative capability of the damaged bone caused by excising the tumour while the patient is undergoing chemotherapy[7]. Our study further corroborates these findings as when BMP-2 is delivered alone, ectopic bone formation is found surrounding the tibia, but when the same hydrogel loaded with the same concentration of BMP-2 was delivered and combined with systemic chemotherapeutics the stimulatory effect was significantly diminished (see **Figure 7**). Interestingly, similar to what we demonstrated using our spheroid model, the long-term effect of the combined therapy (systemic doxorubicin and localised miR-29b) significantly reduced osteolysis due to decreased tumour burden and maintained bone architecture with little to no ectopic bone formation, even in the presence of chemotherapeutics. This anti-cancer and pro-osteogenic effect of miR-29b delivery may extend to other types of cancer. In multiple myeloma, for example, elevated osteoclast activity together with impaired osteoblast function is commonly triggered by tumour cells which proliferate in the bone marrow[15b]. Our miR-29b:pBAE nanoparticles could potentially be delivered to not only treat the tumour cells but also repair osteoblast function. Furthermore, as bone is one of the most common locations of cancer cell metastasis, any therapeutic that inhibits bone tumour growth whilst simultaneously aiding in bone regeneration which the patient is undergoing chemotherapy would significantly benefit not only osteosarcoma patients but any cancer patient with bone metastases (breast, lung, lymphoma, myeloma, prostate and thyroid)[48].

Taken together, this study introduces the therapeutic potential of localised nanoparticle-mediated miR-29b delivery in supressing osteosarcoma growth whilst simultaneously providing the surrounding damaged bone the cues needed to normalise the dysregulation of bone homeostasis caused by tumour growth. A smart delivery vehicle was developed to enable targeted, efficient, local and sustained delivery of miRNAs or growth factors, which can easily be integrated when excising the tumour as a potential add-on to conventional chemotherapy. This novel combined therapy could further improve clinical outcome by significantly reducing primary tumour mass and provide vital information that can inform the design of future combination therapies for these young patients.

## Materials and Methods

### Cell Culture

Human osteosarcoma cell line (SaOS2, HTB-85) and human Mesenchymal Stromal Cells (MSCs, PCS-500-012) were commercially purchased from ATCC. K7 murine osteosarcoma cell line (K7M2-WT, CRL-2836) isolated from the proximal tibia of a BALB/c mouse were commercially purchased from ATCC. Expansion of all cell types was conducted in normoxic conditions and Expansion Medium [Dulbecco’s modified Eagle medium (DMEM; Biosciences) supplemented with 10 % foetal bovine serum (FBS, Gibco), 1% penicillin (100 U/mL) and 1% streptomycin (100 μg/mL) (Biosciences)], was changed twice weekly. Human MSCs and osteosarcoma cells were used at the end of passage 5 for *in vitro* analysis and murine osteosarcoma cells were used at the end of passage 7 for *in vivo* inoculation.

### Synthesis of poly-beta-amino-esters (pBAE) polymers

Synthesis of pBAEs was performed via a two-step procedure, as previously described[27]. First, an acrylate-terminated polymer (termed C6 polymer) was obtained by the addition reaction of 5-amino-1-pentanol (0.426 g, 4.1 mmol) and hexylamine (0.422 g, 4.1 mmol) to 1,4-butanediol diacrylate (2.0 g, 9.1 mmol). The reaction was carried out at 90 °C for 24 h. Second, C6 polymer was end-capped with thiol-terminated arginine peptide (Cys-Arg-Arg-Arg) at a 1:2.1 molar ratio in dimethyl sulfoxide (DMSO) in an overnight, room temperature reaction. The resulting polymer was purified and collected by precipitation in a mixture of diethyl ether and acetone (7:3). pBAE polymer structure was confirmed by ^1^H-NMR (400 MHz Varian (NMR Instruments, Clarendon Hills, IL)).

### miR-29b-pBAE nanoparticle formation and characterization

MiR-29b:pBAE nanoparticles were performed by mixing equal volumes of pBAEs polymer and miR-29b in 12.5 mM acetate buffer (AcONa) at their appropriate concentration. Briefly, pBAE stock solutions in DMSO (100 mg ml−1) were diluted in 12.5 mM AcONa at appropriate concentrations to obtain the desired pBAE:miR-29b weight/weight ratio. The pBAE polymer was added to a solution of miR-29b, incubated at room temperature for at least 5 min, and precipitated in two volumes of PBS 1×. The nanoparticles were purified by centrifugal filtration (10 kDa MWCO Amicon Ultra-4 mL Centrifugal Filters, Millipore), and filtered through a sterile 0.22 μm membrane. The resulting nanoparticles were characterized by an agarose retardation assay and dynamic light scattering (DLS). To assess miR-29b retardation, different miR-29b-to-pBAE (Horizon Discovery, miRIDIAN microRNA Human hsa-miR-29b-3p – Mimic) ratios (w/w) between 10 and 400 were studied. pBAE:miR-29b complexes were freshly prepared and loaded in 2% E-Gel Precast Agarose Gels (Thermo Fisher), run following the manufacturer’s instructions, and visualized in fluorescence mode. Biophysical characterization of nanoparticles was performed using a ZetaSizer Nano ZS equipped with a He–Ne laser (λ 1⁄4 633 nm) at a scattering angle of 137° (Malvern Instruments Ltd, United Kingdom). Hydrodynamic diameter (nm), PDI, and surface charge of nanoparticles were measured.

### Cellular uptake and transfection efficiency

To determine the optimum concentration of pBAE nanoparticles and culture conditions for transfection, human MSCs were seeded onto 24 well plate with polymer cover slip (Corning, P24-1.5P) at a seeding density of 2×10^4^ cells/cm^3^. pBAE:miRNA complexes were performed at a concentration of 100:1 miR29-b:pBAE ratio. After seeding, MSCs were transfected with 10nM, 20nM, or no pBAE nanoparticles loaded with (fresh expansion media) for 4 hours. Following transfection cells were either fixed overnight in 4% paraformaldehyde for confocal imaging, or fresh media was applied (either expansion medium or osteogenic medium [expansion medium + 100nM dexamethasone, 50 mg/mL ascorbic acid, and 10mM β-glycerol phosphate; all Sigma-Aldrich]) and cultured for a further 7 days. After 7 days of culture ALP expression into the media was used as measure of transfection efficiency. This was measured using a colorimetric assay of enzyme activity (SIGMAFAST p-NPP Kit; Sigma Aldrich), which uses p-nitrophenyl phosphate (pNPP) as a phosphatase substrate with ALP enzyme (Sigma Aldrich) as a standard[49].

To validate the transfection efficiency of the pBAE nanoparticles the transfection potential was directly compared to the commercially available transfection reagent Lipofectamine (RNAiMAX, Thermofisher). Briefly, human MSCs were seeded at 2×10^4^ cells/cm^2^ in a standard 24 well plate. After seeding MSCs were transfected with 20nM of either pBAE:miR-29b complexes for 4 hours or 20nM Lipfactamine:miR-29b complexes for 24 hours. Following transfection fresh media was applied (either expansion medium or osteogenic medium) and cultured for a further 7 days. ALP activity was measured after 7 days of culture.

To validate the pro-osteogenic, anti-angiogenic potential of miR-29b delivery, human MSCs were seeded at 2×10^4^ cells/cm^2^ in a standard 24 well plate. Following seeding MSCs were transfected with 20nM of either pBAE:miR-29b (*miR-29b*) complexes or pBAE:siRNA (*Scramble*) complexes (Qiagen, AllStars Negative Control siRNA) or cultured in standard control medium (*control*) for 4 hours. Following transfection fresh osteogenic media was applied and the cells were cultured for a further 7 days. ALP expression was measured after 7 days of culture. An enzyme-linked immunosorbent assay was used to quantify the levels of VEGF (Bio-Techne, MN, USA) released by the cells. The cell culture media was analysed following 7 days of culture. Assays were carried out as per the manufacturer’s protocol and analysed on a microplate reader at a wavelength of 450 nm as previously described[7, 35a, 50]. To assess DNA content, cells were subjected to cell lysis by three cycles of freeze–thawing in molecular grade water[51]. DNA content was measured using Quant-iT PicoGreen dsDNA Assay (BD Biosciences), with calf thymus DNA as a standard as previously described[7, 52].

To validate the apoptotic nature of miR-29b delivery to cancer cells human osteosarcoma cells (SaOS2) and mouse osteosarcoma cells (K7M2) were seeded onto 24 well plate with polymer cover slip (Corning, P24-1.5P) at a seeding density of 2×10^4^ cells/cm^3^. pBAE:miR-29b (*miR-29b*) and pBAE:siRNA (*Scramble*) complexes were prepared as mentioned above and following seeding both cell types were transfected with 20nM of either complex or cultured in osteogenic medium (*control*) for 4 hours. After transfection, cells were either fixed overnight in 4% paraformaldehyde for confocal imaging, or fresh media was applied and cultured for a further 3 days. ALP expression into the media was assessed after 3 days of culture. DNA content was also measured as mentioned above following 3 days of culture.

### Osteosarcoma Spheroid Model

Microwells were created using moulding system as previously described[7, 53]. Briefly, sterile molten agarose solution (4% w/v) was pipetted into a well of a 6 well plate, where custom made 3D printed positive moulds were inserted. Once cooled, the positive mould was pulled from the agarose, leaving 401 microwells within each well. All agarose microwells were soaked overnight in DMEM prior to cell seeding. A 3:1 seeding ratio of human MSCs to osteosarcoma cells (MSC:SaOS2) was used as previously described[7]. Cells were seeded into the microwells by pipetting an appropriate density (4000 cells/microwell) into each well. After seeding, plates were centrifuged at 700 × g for 5 min to collect cells at the bottom of each well. Plates were then incubated in expansion medium for 72 h to allow for formation of tumour spheroids. pBAE:miR-29b nanoparticles were prepared as mentioned above and the osteosarcoma tumour spheroids were either transfected with 20nM concentration of pBAE nanoparticles for 4 hours or cultured in expansion medium for 4 hours. Following transfection fresh media was applied and the tumour spheroids were cultured under the following culture conditions for a further 7 days of culture: [1] *Control* – cultured in osteogenic medium without transfection; [2] *Control+Dox* – cultured in osteogenic medium plus 1.8 μM of Doxorubicin (IC_50_ concentration previously determined[7]) without transfection; [3] *miR-29b* – cultured in osteogenic medium and 4 hour miR-29b transfection; [4] *miR-29b + Dox* – cultured in osteogenic medium plus 1.8 μM of Doxorubicin and 4 hour miR-29b transfection.

After 1 week, tumour spheroids were liberated from the microwells as previously described[7]. Spheroids were prepared in one of the following ways: 1) snap-frozen and stored at −80 °C for DNA analysis; 2) harvested and aliquoted for flow cytometry; 3) snap-frozen using liquid nitrogen and stored at −80 °C for PCR analysis; or 4) embedded within 2% agarose and fixed overnight in 4% paraformaldehyde before being placed in PBS and refrigerated for histochemical analysis.

### DNA Analysis

To assess DNA content for all constructs, 0.5 mL of papain digestion buffer (100 mm sodium phosphate buffer containing 10 mm L-cysteine [Sigma-Aldrich], 125 μg mL−1 papain [Sigma-Aldrich], and 5mmNa2EDTA [Sigma-Aldrich] in ddH2O at pH 6.5) was added to the spheroids that were placed on a rotator in an oven at 60 °C overnight, as previously described[7]. Once the spheroids were digested, DNA content was performed using Quant-iT PicoGreen dsDNA Assay.

### Calcium Deposition

Calcium deposition within the tumour spheroids was measured using the Calcium LiquiColor Test (Stanbio Laboratories) according to the manufacturer’s protocol. Briefly, a cell lysate was prepared by digesting the spheroids in a 0.5M hydrochloric acid solution at 60 °C overnight, as previously described[7, 52].

### PCR Analysis

RNA from spheroids was extracted using Trizol Reagent and assessed for concentration and purity using the NanoDrop 2000c UVis spectrophotometer. RNA was equalized and reverse transcribed using the Applied Biosystems High-Capacity cDNA reverse transcription kit. Real-time PCR was carried out on triplicate cDNA samples with the use of the CFX96 Touch Real-Time PCR Detection System (Bio-Rad Laboratories, California). Real-time PCR for the detection of *MMP2, MMP9, β-Catenin, Bcl-2, Casp 9, Casp 3*, and *BAX* mRNA was performed using the TaqMan fast universal PCR Master Mix (Applied Biosystems) and predesigned TaqMan gene expression primers. mRNA amounts were normalized relative to the housekeeping gene Ribosomal Protein 18s. For the assessment of growth factor gene expression Sigma primers for *VEGF, RUNX2, Col1A, TGFβ3*, and *ALP* (Table S1, Supporting Information) were used, and mRNA amounts were normalized to Glyceraldehyde 3-phosphate dehydrogenase (GAPDH) as housekeeping gene.

### Flow Cytometry

Prior to cell seeding MSCs and SaOS2 cells were stained with cell tracker dyes; QTracker705 and QTracker585 (Thermo Fisher), respectively, as previously described[7]. Cells were seeded and transfected and cultured for a further 7 further days under their respective culture conditions. Following 1 week of cultures, spheroids were harvested and aliquoted into triplicates such that there was a minimum of 100 spheroids per sample. A single cell suspension was created by incubating the tumour spheroids with 0.25% Trypsin-EDTA at 37 °C for 15 min. Following 1 wash with PBS, cells were stained with LIVE/DEAD Fixable Near-IR fluorescent dye (L10119, Thermo Fisher) according to the manufacturer’s instructions. Following fixation, the cell suspensions were analysed by flow cytometry using a BD LSR II HTS-1 flow cytometer (BD Biosciences) to determine the ratio of MSC+ to SaOS2+ cells for each culture condition. All data was analysed using Flowjo software (Flowjo LLC), gating strategy can be found in **Supplementary Fig. 3**.

### Histology

Following embedding within 2% agarose and fixation overnight samples were dehydrated and embedded in paraffin using an automatic tissue processor (ASP300; Leica, London, UK). All samples were sectioned at a thickness of 6 μm using a rotary microtome (Leica). Sections were stained with H&E and 2% Alizarin Red solution (all Sigma-Aldrich).

### Preparation of Transfected Wild Type K7M2 Cells

K7M2 WT cells were seeded into a 96 Well White/Clear Bottom Plate (Thermo Fisher, 165306) at a seeding density of 2×10^4^ cells in 100µL per well. To determine the optimum transfection ratio multiple complexes were created (1.5:1, 3:1, 6:1) using ViaFect Transfection reagent (Promega, E4981) and pGL4.51 [luc2/CMV/Neo] plasmid vector (Promega, E1320) as per the manufacturer’s specifications and incubated with K7M2 cells for 24 hours. Following 24 hours of incubation the cells were washed twice with PBS and fresh medium was added to the cells after which they were further cultured for another 48 hours. The selective drug Geneticin-418 (Thermo Fisher, 10131035) was then added at a concentration of 200 µg/mL to kill non-transfected cells. To validate transfection and determine optimum transfection ratio *in vitro* bioluminescence imaging was performed. Briefly, a 200X Luciferin stock solution (30mg/mL) was prepared in sterile water. From that stock solution a 150µg/mL working solution was prepared in expansion medium. The working solution was added directly onto the transfected K7M2 cells and bioluminescence was measured using Infinite M Plex (Tecan) plate reader every 10 mins, up to 40 mins to determine the kinetic curve and find the peak imaging timepoint for the cells (see **Supplementary Fig. 4 (A)**).

For *in vivo* inoculation transfected K7M2 osteosarcoma cells were prepared by seeding 6×10^5^ cells into each well of a six-well plate. A 6:1 transfection ratio complex was created as was deemed the optimum transfection ratio and the K7M2 cells transfected with the luciferase reporter as described above.

To assess the stability of the luciferase reporter *in vitro* bioluminescence imaging was performed on the transfected K7M2 cells for multiple cell passages (see **Supplementary Fig. 4 (B)**). Passage two transfected cells were used for orthotopic implantation.

### Development of metastatic osteosarcoma murine model

All mouse procedures were conducted at the Koch Institute for Integrative Cancer Research at the Massachusetts Institute of Technology (MIT) under the protocol approved for this study by the Institutional Animal Care and Use Committee (IACUC). To assess the optimum concentration of osteosarcoma cells needed to induce tumour formation, two concentrations of transfected K7M2 cells were evaluated 0.3×10^6^ cells and 1×10^6^ cells[30]. Six female BALB/c mice (3 *0.3×10^6^ cells*, 3 *1×10^6^ cells*) aged 6 weeks underwent inoculation with either of the cell concentrations (see **Supplementary Figure 5**). Briefly, the BALB/c mice were anesthetized using 2% isoflurane. Analgesia, slow-release buprenorphine (1mg/kg) was administered subcutaneously. A small incision into the skin above the tibial plateau was made and the muscle was dissected away. A small hole was created on the tibial plateau into the medullary canal using a 26G needle. 10 µL of the luciferase labelled K7M2 cells at either concentration was injected into the proximal tibia using a 27G Hamilton syringe.

### Synthesis of amino-modified Hyaluronic Acid (HA-SS-NH_2_)

Synthesis of HA-SS-NH_2_ was performed as previously described[54]. Briefly, 60 kDa-sodium hyaluronate (1% w/v in MES buffer) was modified with Cysteamine Dihydrochloride through N-(3-(dimethylamino)propyl)carbodiimide (EDC) and N-hydroxysuccinimide (NHS) chemistry. The reaction was performed at room temperature for 12 h. The product HA-SS-NH2 was purified by dialysis, freeze-dried, and stored at −20 °C. HA-SS-NH2 polymer structure was confirmed by 1H-NMR (400 MHz Varian (NMR Instruments, Clarendon Hills, IL)).

### Development of Hyaluronic Acid (HA) injectable delivery system

Amine-modified Hyaluronic Acid was dissolved in phosphate buffer (pH = 7.4) containing the miR-29b-pBAE nanoparticles to obtain a 10 % (w/v) HA solution. 8-arm-PEG-NHS crosslinker was also dissolved in phosphate buffer (pH = 7.4) to obtain a 10 % (w/v) solution. For *in vitro*, the two polymer solutions were vigorously mixed together for 10s inside cylindrical plastic moulds (diameter: 5.00 mm; height: 2.50 mm). Hydrogel disks were allowed to react for 5 min to ensure full gelation. For *in vivo*, the two polymer solutions were loaded in a double-channel syringe coupled to a mixer allowing the hydrogel formation once is injected.

### Surgical Procedure

Forty female BALB/c mice aged 6 weeks underwent inoculation with 0.3×10^6^ K7M2-Luc cells according to the same protocol as mentioned above. The average mouse weight was 18.5g at the time of inoculation. Palpable tumours were allowed to form two weeks and once a suitable sized tumour has formed, the animals (*n*=8) were randomly assigned into different treatment groups: [1] ***No Treatment***: Tumour cell inoculation with no further treatment; [2] ***Dox Alone:*** Tumour cell inoculation plus tail vein (i.v.) injection of Doxorubicin (Fisher, BP2516-10) 2mg/kg biweekly for three weeks; [3] ***Hydrogel Alone:*** mice were anesthetized with 1–2% isoflurane and non-invasively 100 µL of HA hydrogel was injected around the tumour site; [4] ***miR-29b:*** mice were anesthetized with 1–2% isoflurane and non-invasively 100 µL of HA hydrogel loaded with fluorescently-labelled (640 ex, 680 em) pBAE:miR-29b nanoparticles (750µg/kg miR-29b) was injected around the tumour site; [5] ***miR-29b + Dox:*** mice were anesthetized with 1–2% isoflurane and non-invasively 100 µL of HA hydrogel loaded with fluorescently-labelled pBAE:miR-29b nanoparticles was injected around the tumour site. Following hydrogel implantation mice were treated with i.v. injection of Doxorubicin (2mg/kg) biweekly for three weeks. Thirty-one out of forty mice grew primary tumours.

The tumour size was measured twice a week via calliper measurements, and the tumour volume was calculated using the following formulas: leg volume = length x (width)^2^x 0.5; tumour volume = leg Volume on day X /leg Volume on day 0. Body weight was measured contemporaneously with tumour volume. Mice were euthanized when tumours reached a volume of 1000 mm^3^or for otherwise poor body condition. Nine mice were removed from the study as no bioluminescent signal was recorded and no palpable tumour formed in these mice in the 45 days following tumour cell inoculation.

### Bioluminescent (BLI) Imaging

Briefly, mice were shaved then anesthetized with 1–2% isoflurane using a calibrated vaporizer inside an IVIS Spectrum (PerkinElmer). Non-invasive longitudinal monitoring of tumour progression and nanoparticle release was followed by scanning mice with the IVIS Spectrum-bioluminescent and fluorescent imaging system (PerkinElmer). Luciferin (150mg/kg) was injected intra-peritoneally and at each timepoint a kinetic curve was performed to determine the peak signal time (∼15-20 mins). Whole-animal bioluminescent and fluorescent imaging was performed biweekly for 18 days following treatment.

### X-Ray Microtomography (µCT)

Micro-CT images of the tibias was performed in Skyscan 1276 (Bruker) µCT pre-treatment, 2- and 4-weeks post-treatment to monitor osteolysis and metastasis. Briefly, mice were anesthetized with 1–2% isoflurane using a calibrated vaporizer inside a Skyscan 1276 µCT and noninvasively imaged. Scans were performed using a voxel resolution of 16 µm. Data was analysed using ImageJ BoneJ plugin.

### Histological Analysis

All of the tibias were removed from the mice at their respective end points and fixed overnight at 4°C in 10% formalin. Tibias were also decalcified for 1 week before tissue processing. All samples were dehydrated and embedded in paraffin using an automatic tissue processor (Leica ASP300, Leica). All samples were sectioned with a thickness of 8 µm using a rotary microtome (Leica Microtome RM2235, Leica). Sections were stained with H&E to assess tumour growth or Goldner’s Trichrome to assess bone regeneration.

### BMP-2 Delivery

To validate the clinical relevance of localised delivery of miR-29b to induce bone remodelling within the bone and normalise the dysregulation of bone homeostasis caused by the tumour and chemotherapeutics, the bone formation potential was directly compared to BMP-2 delivery which is known to induce bone formation. Sixteen BALB/c mice aged 6 weeks underwent inoculation with 0.3×10^6^ K7M2 cells according to the same protocol as mentioned above. Once a suitable sized tumour has formed, the animals (*n*=8) were randomly assigned into two different treatment groups: [1] ***BMP-2:*** mice were anesthetized with 1–2% isoflurane and non-invasively 100 µL of HA hydrogel loaded with 5µg BMP-2[35] (Peprotech, 120-02) was injected around the tumour site; [2] ***BMP-2 + Dox:*** mice were anesthetized with 1–2% isoflurane and non-invasively 100 µL of HA hydrogel loaded with was injected around the tumour site. Following hydrogel implantation mice were treated with i.v. injection of Doxorubicin (2mg/kg) biweekly for three weeks. Tumour volume and body weight were measured every other day. Six mice were removed from the study as no bioluminescent signal was recorded and no palpable tumour formed in these mice in the 45 days following tumour cell inoculation.

### BMP-2 enzyme-linked immunosorbent assay (ELISA)

For *in vitro* BMP-2 release studies, the two polymer solutions (loaded with 5µg BMP-2) were vigorously mixed together for 10s inside cylindrical plastic moulds (diameter: 5.00 mm; height: 2.50 mm). Hydrogel disks were allowed to react for 5 min to ensure full gelation. The hydrogels were then culture in expansion media for 35 days in culture. A BMP-2 ELISA (R&D systems) was used to quantify the levels of BMP-2 released by the hydrogels. The cell culture media were analysed at the specific time points detailed above. Assays were carried out per the manufacturer’s protocol and analysed on a microplate reader at a wavelength of 450 nm.

### Statistical Analysis

Results were expressed as mean ± standard deviation. Statistical analysis was performed using the following variables: (1) when there was two groups and one time-point a standard two-tailed t-test was performed, (2) when there were more than two groups and one time-point a one-way analysis of variance (ANOVA) was performed, and (3) when there were more than two groups and multiple time points a two-way ANOVA was performed. For *in vivo* studies multiple comparisons among groups were determined using Kruskal–Wallis test with uncorrected Dunn’s test. Kaplan–Meier survival curve statistical analysis was determined using the two-tailed Mantel–Cox test. No specific pre-processing of data was performed prior to statistical analyses. All analyses were performed using GraphPad (GraphPad Software, La Jolla California USA, www.graphpad.com). For all comparisons, the level of significance was p ≤ 0.05.

## Acknowledgments

The authors thank the Division of Comparative Medicine at MIT for the assistance with animal housing. The authors thank Swanson Biotechnology Center at the Koch Institute for Integrative Cancer Research at the Massachusetts Institute of Technology (MIT) for assistance with animal experiments and facilities, especially the microscopy, preclinical imaging & testing, flow cytometry, and histology cores. The authors thank Dr A.M. Hayward for clinical assistance in developing the osteosarcoma murine model. The authors thank G. Paradis for FACS assistance with Cancer Center Support (FACS core). The authors thank Kathleen S. Cormier for histology assistance. The authors thank M. Cornwall-Brady for assistance with µCT scanning and analysis. The authors thank J. Kuhn for confocal imaging assistance.

## Funding

This project has received funding from the European Union’s Horizon 2020 research and innovation programme under the Marie Skłodowska-Curie Grant Agreement No. 839150 and the Gillian Reny Stepping Strong Center for Trauma Innovation.

## Author contributions

F.E.F was responsible for technical design, development of osteosarcoma tumour model, all in vitro pre-screening experiments, developing the osteosarcoma animal model, performing all of the animal surgeries, all IVIS imaging and CT scans, data interpretation, histological analysis and drafting the paper. P.D.P assisted with the animal surgeries and developed and characterised the pBAE nanoparticles and HA injectable delivery system. C.R.R.J assisted with the characterisation and in vitro release profile of the pBAE nanoparticles from the HA hydrogel. O.M helped with all of the PCR analysis. D.J.K. and N.A conceived and helped design the experiments, oversaw the collection of results and data interpretation, and finalized the paper.

## Competing interests

Research undertaken in Daniel Kelly’s laboratory at Trinity College Dublin is part-funded by Johnson & Johnson. The authors declare no other competing interests.

## Data and materials availability

All data needed to evaluate the conclusions in the paper are present in the paper and/or the Supplementary Materials. Additional data related to this paper may be requested from the authors.

## Supplementary Information

### Determination of the optimum transfection conditions

To validate the optimum concentration of nanoparticles for transfection a 10nM and 20nM concentration[55] was evaluated. First, we studied the uptake of the pBAE nanoparticles (pink; AlexaFluor 647) by the cells using confocal imaging. Both concentrations of pBAEs were taken up by the cells, however there was higher uptake when transfected with the 20nM concentration (see Supplementary Fig. 6 (A)). To determine the optimum culture conditions to promote osteogenesis in MSCs, following a 4-hour transfection period with pBAE:miR-29b complexes, ALP activity was evaluated after a further 7-day culture period in either expansion medium or osteogenic medium. As expected, culturing MSCs in osteogenic medium versus expansion medium, regardless of transfection, led to a significant increase in ALP activity when directly compared to its expansion medium control (see Supplementary Fig. 6 (B)). Interestingly, transfection with miR-29b (regardless of culture condition) led to a significant increase in ALP activity compared to the non-transfected control. Therefore it was determined that a culture condition of 20nM pBAE concentration plus further culture in osteogenic medium will hereafter be used for all in vitro studies, as it induced the highest ALP activity following transfection.

To validate the transfection efficiency of the pBAE nanoparticles, the transfection potential was directly compared to the commercially available transfection reagent Lipofectamine. Here, it is clear that the pBAE delivery vector offers superior transfection capabilities, as there was significantly higher ALP activity, thereby confirming miR-29b functionality, in MSCs transfected with pBAE nanoparticles compared to its counterpart transfected using Lipofectamine (see Supplementary Fig. 6 (C)).

### Development of the metastatic orthotopic osteosarcoma mouse model

To allow for continuous monitoring of tumour growth over time using *In Vivo* Imaging System (IVIS), K7M2 cells were transfected with Luciferin plasmid vector using ViaFect Transfection reagent (6:1 Transfection Reagent: DNA ratio) which was deemed the optimum ratio for transfection (see **Supplementary Fig. 6 (A)**). Stability of the transfected cell line was then monitored over 7 passages of cells. There was a significant difference in luminescence from passage 1 to passage 2 (3295.5 p/sec vs 356.67 p/sec) (see **Supplementary Fig. 6 (B)**). However, there was no significant difference in luminescence of the transfected cells at passage 2 and passage 7 (356.67 p/sec vs 320.67 p/sec). Linear regression modelling (passage 2 – passage 7) did not reveal significant differences in radiance of the transfected cells over time (R^2^= 0.016, p =0.81). With this in mind, passage 2 K7M2 transfected cells were used for all *in vivo* inoculation studies.

For the development of the osteosarcoma model six female BALB/c mice were inoculated (6 left limbs) with K7M2-Luc cells at two different concentrations 1×10^6^ (*n* = 3) as previously described^[30]^, and 0.3×10^6^ cells (*n* = 3). The average mouse weight was 18.3g at the time of inoculation. All six grew primary tumours with the 1×10^6^ group growing at an average of 15 days and the 0.3×10^6^ group growing at an average of 14 days. There was no significant difference in luminescence at each time point between the 0.3×10^6^ and the 1×10^6^ groups (see **Supplementary Fig. 5 (A)**). There was a significant increase in luminescence over time in both groups, as the tumours grew. MicroCT analysis three weeks after inoculation showed no significant difference in BV/TV between the two groups, but there was a trend towards a significant decrease (p = 0.08) in BV/TV in the 0.3×10^6^ group. Furthermore, 3D reconstructions revealed a trend towards increased osteolysis due to tumour growth in the 0.3×10^6^ group (see **Supplementary Fig. 5 (B)**). This was further verified with histological analysis, as the 1×10^6^ group still had some trabecular bone present within the marrow cavity (denoted by black arrow heads) whereas the 0.3×10^6^ group had little to no trabecular bone present due to extensive tumour growth (see **Supplementary Fig. 5 (C)**). With the slight increase in palpable tumour growth (∼14 days), increased osteolysis as seen in the µCT reconstruction images, and the trend towards increased luminescent signal, 0.3×10^6^cells was deemed the optimum cell concentration to induce osteosarcoma tumour growth.

---

**Supplementary Fig. 1.**
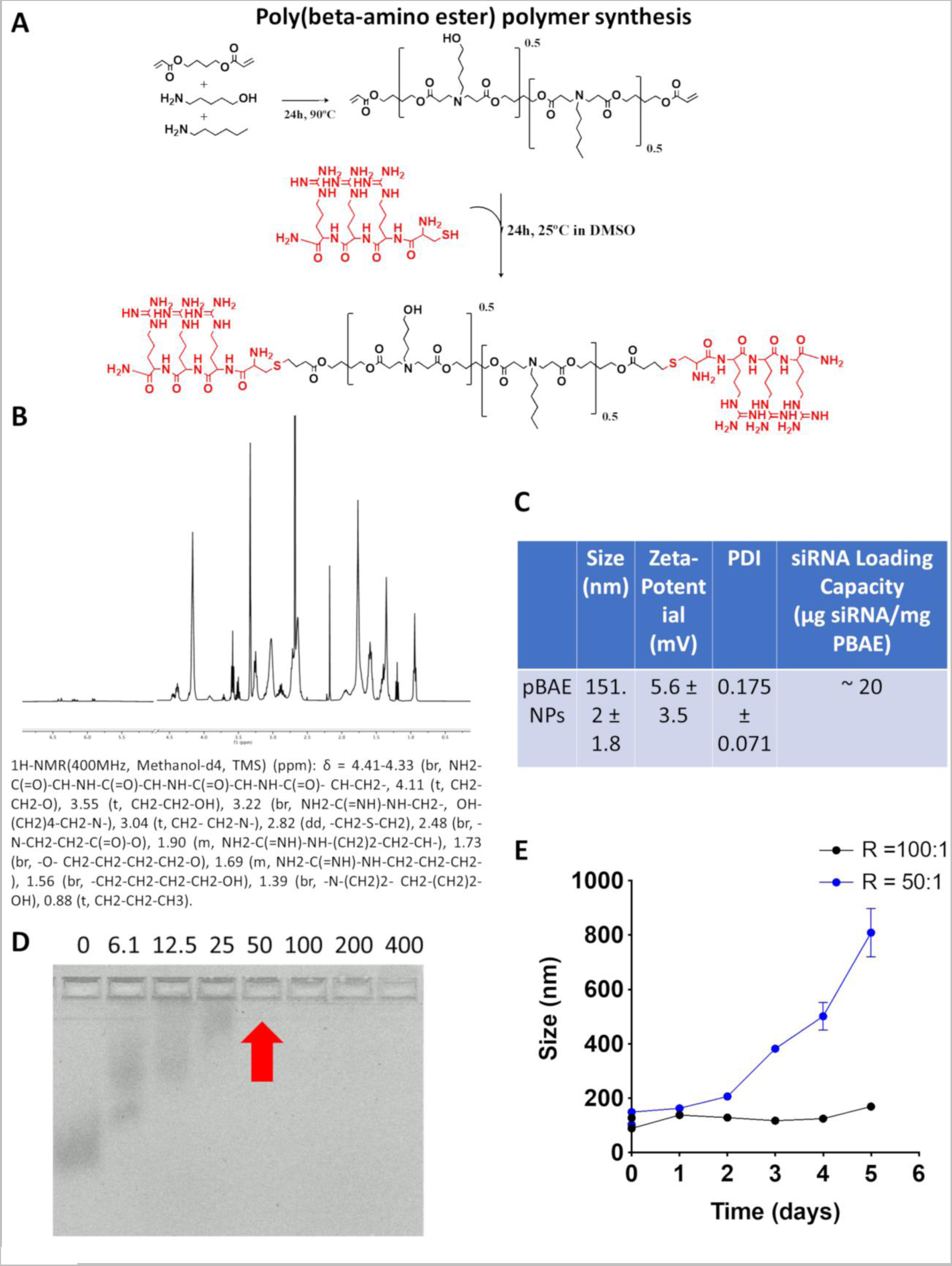
Synthesis and Characterisation of pBAE nanoparticles. **(A)** Synthesis of arginine modified poly (beta-amino) formulation was performed via a two-step procedure, First, a mixture of 5-amino-1-pentanol, hexylamine, and 1,4-butanediol diacrylate (0.5:0.5:1.2) were used for the synthesis of pBAE C6 polymer. Then, the arginine-modified pBAE (C6-CR3) was formulated by mixing acrylate-terminate pBAE polymer with polyarginine peptide containing a cysteine amino acid (Cys-Arg-Arg-Arg). **(B)** 1H-NMR characterization of arginine-modified poly(beta-amino ester) (C6-CR3) polymer. **(C)** Biophysical characterization of miR-29b NPs was determined by Dynamic Light Scattering (DLS). **(D)** Agarose retardation assay of C6-CR3. Nanoparticles were formed using miR-29b and C6-CR3 at different w/w ratios and loaded onto an agarose gel to assess miR-29b mobility by electrophoresis. **(E)** Stability study of miR-29b NPs was determined by DLS.

**Supplementary Fig. 2:**
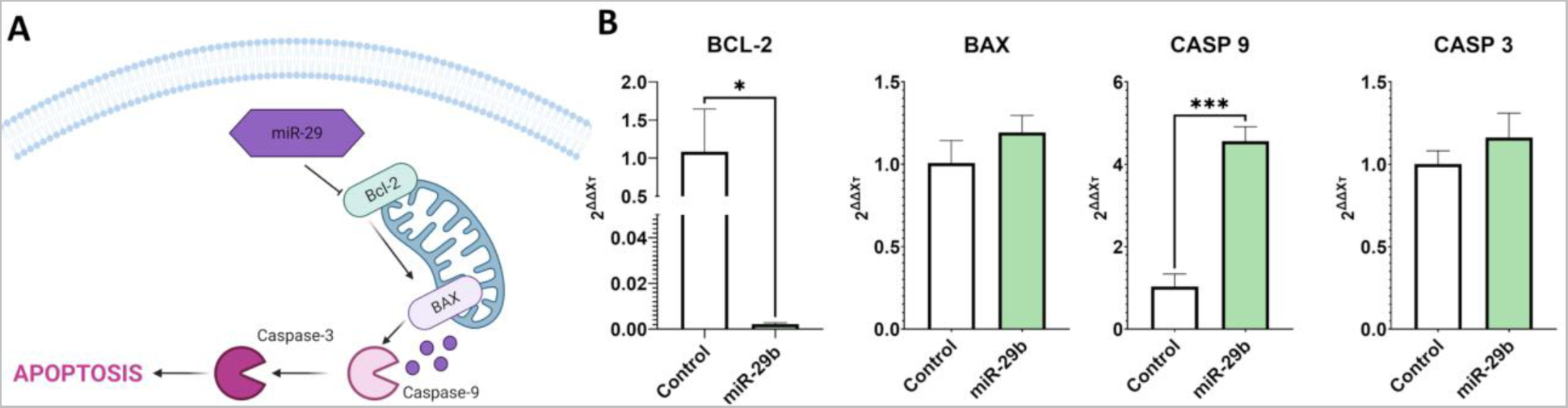
Potential Mechanism behind miR-29b-mediated tumour cell apoptosis. **(A)** Schematic of the miR-29b/BCL-2 pathway. **(B)** mRNA levels of BCL-2, BAX, Caspase 9 and Caspase 3 were analysed of SaOS2 cells by qRT-PCR. All data was obtained following a 7-day culture period post-transfection and is represented as means ± SD n = 3 (100 spheroids per experimental replicate). Statistical differences were assessed using a student t-test. *p < 0.05, ***p < 0.001.

**Supplementary Fig. 3:**
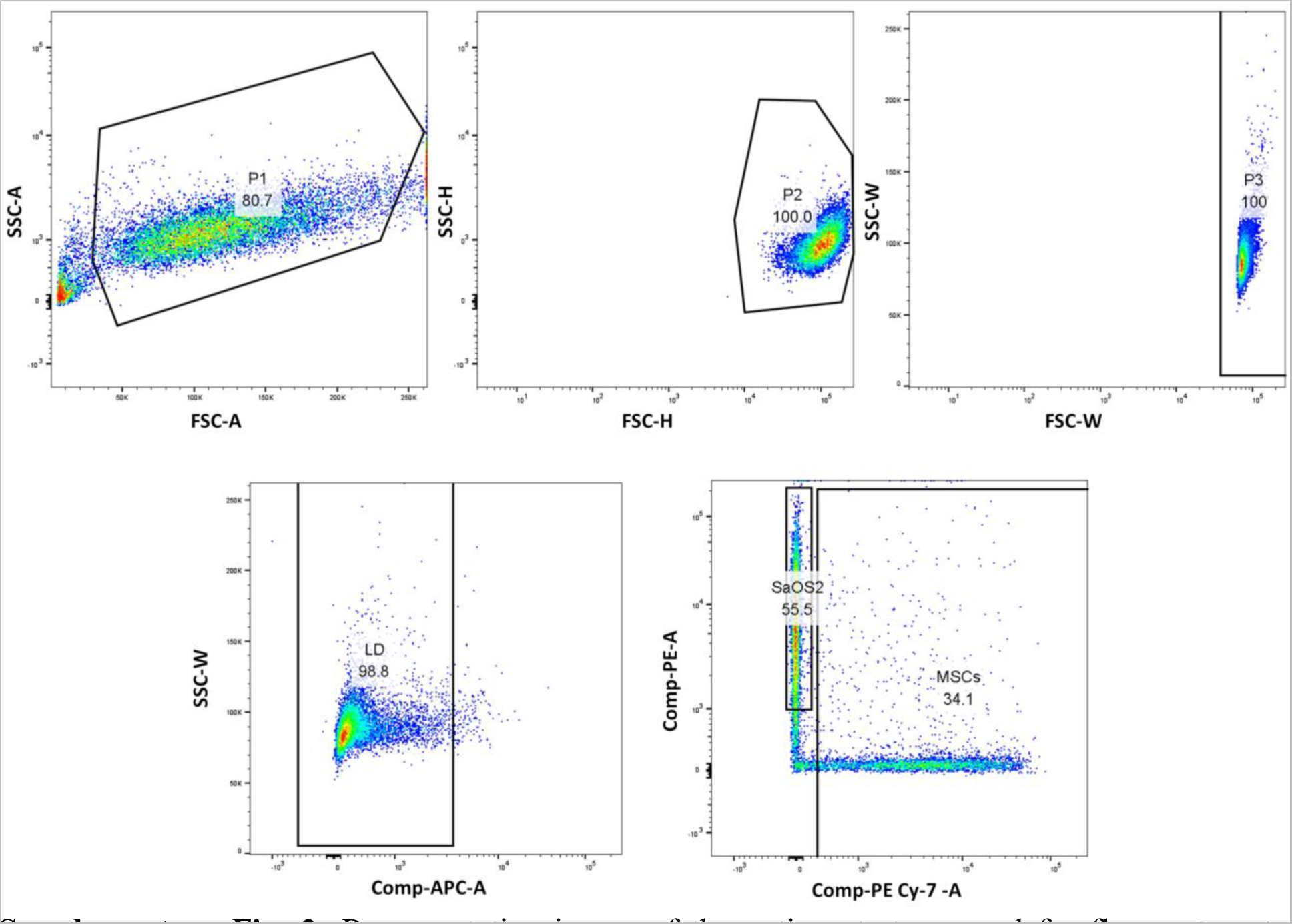
Representative image of the gating strategy used for flow cytometry analysis.

**Supplementary Fig. 4:**
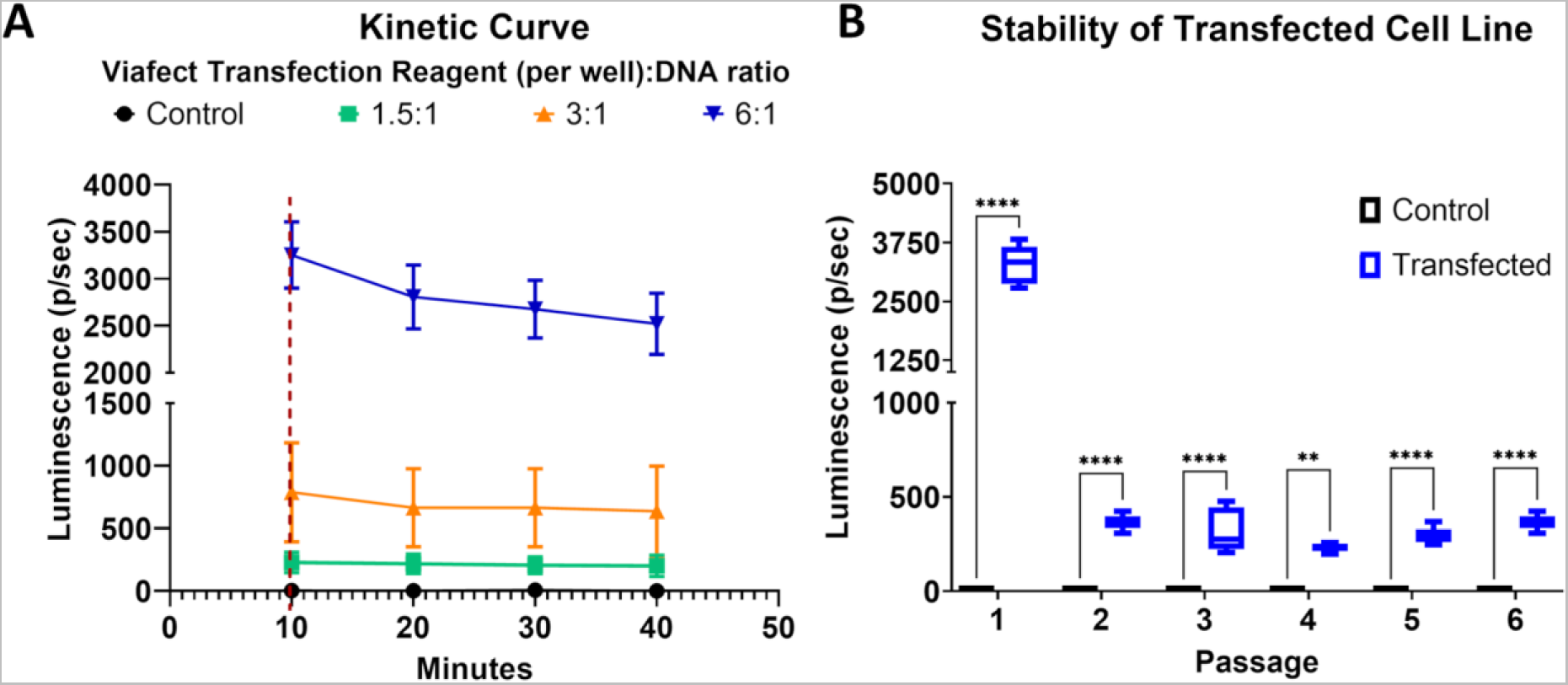
Transfection of K7M2 cells with Luciferin. **(A)** Determining the optimum transection ratio to transfect K7M2 with Luciferase plasmid vector. **(B)** Stability of transfected K7M2 over multiple passaged in vitro. All data is represented as Means ± SD n=6. Statistical differences were assessed using one-way ANOVA. *p < 0.05, **p < 0.01, ***p < 0.001, ****p < 0.0001.

**Supplementary Fig. 5:**
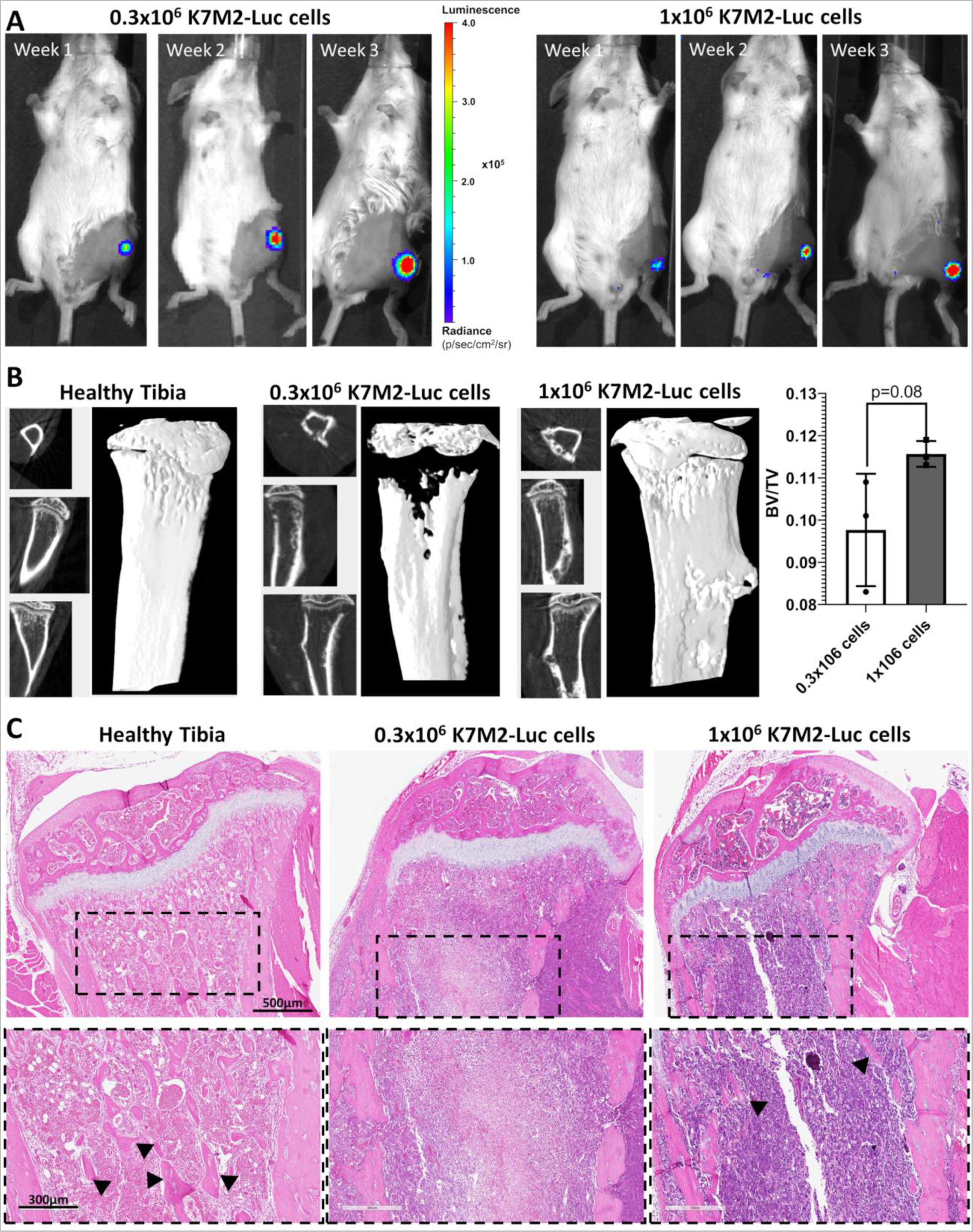
Development of metastatic orthotopic osteosarcoma mouse model. **(A)** One week after injection of K7M2-Luc cells (0.3 or 1 x 10^6^ cells) the signal from labelled cells was localized in the injection site of the right knee of the mouse. On day 21 from inoculation, tumour growth at the injection site (right knee) was macroscopically detectable. **(B)** CT analysis including 3D reconstructions and Bone Volume/Total Volume (BV/TV) following 3 weeks of induction. **(C)** H&E staining revealed densely packed marrow with malignant cells (purple staining) with full trabeculae regression including invasion of the cortical bone and surrounding muscle (Magnification 20X). Black arrow heads denote remaining trabeculae within the marrow cavity.

**Supplementary Fig. 6:**
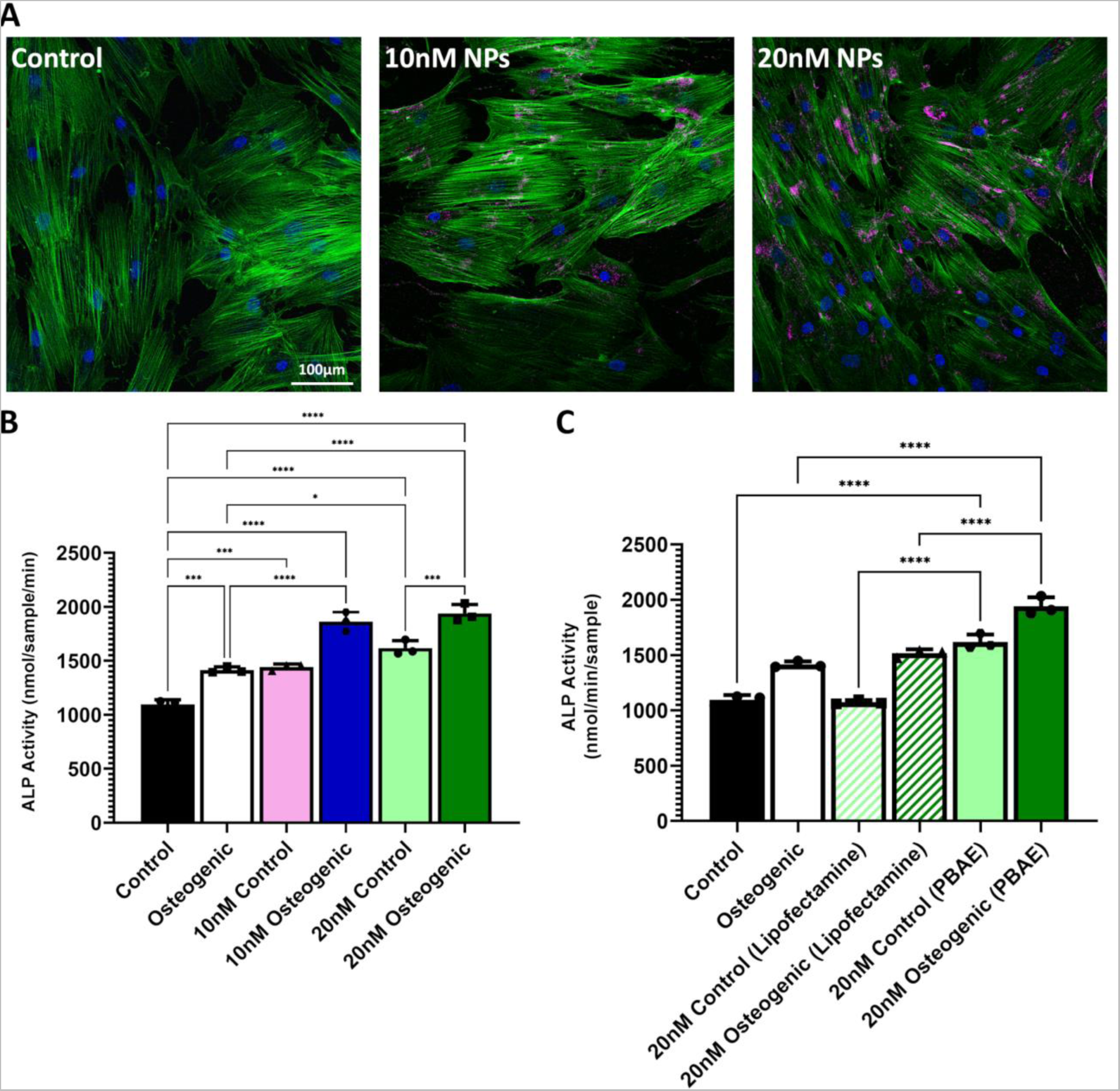
Screening of optimum nanoparticle concentration, *in vitro* culture condition, and nanoparticle for miR-29b delivery. **(A)** Confocal images of human MSCs following 4-hour transfection with fluorescently labelled-pBAE nanoparticles at varying 63 concentrations; Blue (nuclei), Green (actin-cytoskeleton), and pink (mir-29b pBAE nanoparticles). **(B)** ALP activity over 7 days following transfection with miR-29b loaded pBAE nanoparticles at varying concentration (10nM vs. 20nM) cultured in either expansion media (control) or osteogenic culture media (osteogenic). **(C)** ALP activity and over 7 days following transfection with miR-29b using either pBAE nanoparticles or commercially available transfection reagent Lipofectamine 5 cultured in either expansion media (control) or osteogenic culture media (osteogenic). All data is represented as Means ± SD *n*=4. Statistical differences were assessed using one-way ANOVA. *p < 0.05, **p < 0.01, ***p < 0.001, ****p < 0.0001.

**Supplementary Table 1:**
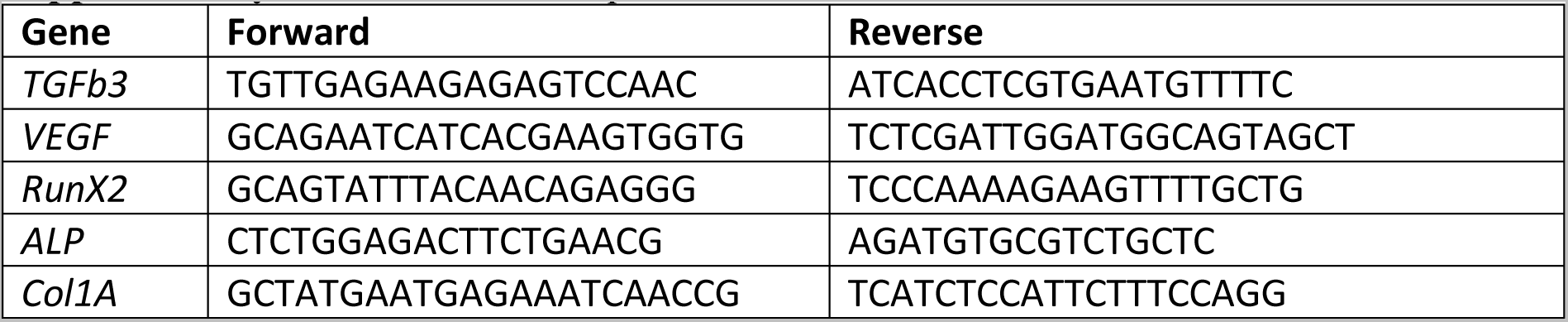
Primer sequences

## Notes

### Competing Interest Statement

The authors have declared no competing interest.

## References and Notes

[1] A. Misaghi, A. Goldin, M. Awad, A. A. Kulidjian, SICOT J 2018, 4, 12, https://doi.org/10.1051/sicotj/2017028.

[2] K. Enterprises, The Ortho Factbook, Knowledge Enterprises Inc, 2001.

[3] a) T. M. Fan, R. D. Roberts, M. M. Lizardo, Front Oncol 2020, 10, 13, https://doi.org/10.3389/fonc.2020.00013; b) K. Winkler, G. Beron, G. Delling, U. Heise, H. Kabisch, C. Purfürst, J. Berger, J. Ritter, H. Jürgens, V. Gerein, Journal of Clinical Oncology 1988, 6 (2), 329, https://doi.org/10.1200/jco.1988.6.2.329; c) N. M. Marina, S. Smeland, S. S. Bielack, M. Bernstein, G. Jovic, M. D. Krailo, J. M. Hook, C. Arndt, H. van den Berg, B. Brennan, B. Brichard, K. L. B. Brown, T. Butterfass-Bahloul, G. Calaminus, H. E. Daldrup-Link, M. Eriksson, M. C. Gebhardt, H. Gelderblom, J. Gerss, R. Goldsby, A. Goorin, R. Gorlick, H. E. Grier, J. P. Hale, K. S. Hall, J. Hardes, D. S. Hawkins, K. Helmke, P. C. W. Hogendoorn, M. S. Isakoff, K. A. Janeway, H. Jürgens, L. Kager, T. Kühne, C. C. Lau, P. J. Leavey, S. L. Lessnick, L. Mascarenhas, P. A. Meyers, H. Mottl, M. Nathrath, Z. Papai, R. L. Randall, P. Reichardt, M. Renard, A. A. Safwat, C. L. Schwartz, M. C. G. Stevens, S. J. Strauss, L. Teot, M. Werner, M. R. Sydes, J. S. Whelan, Lancet Oncol 2016, 17 (10), 1396, https://doi.org/10.1016/s1470-2045(16)30214-5; d) S. S. Bielack, S. Smeland, J. S. Whelan, N. Marina, G. Jovic, J. M. Hook, M. D. Krailo, M. Gebhardt, Z. Pápai, J. Meyer, H. Nadel, R. L. Randall, C. Deffenbaugh, R. Nagarajan, B. Brennan, G. D. Letson, L. A. Teot, A. Goorin, D. Baumhoer, L. Kager, M. Werner, C. C. Lau, K. Sundby Hall, H. Gelderblom, P. Meyers, R. Gorlick, R. Windhager, K. Helmke, M. Eriksson, P. M. Hoogerbrugge, P. Schomberg, P. U. Tunn, T. Kühne, H. Jürgens, H. van den Berg, T. Böhling, S. Picton, M. Renard, P. Reichardt, J. Gerss, T. Butterfass-Bahloul, C. Morris, P. C. Hogendoorn, B. Seddon, G. Calaminus, M. Michelagnoli, C. Dhooge, M. R. Sydes, M. Bernstein, J Clin Oncol 2015, 33 (20), 2279, https://doi.org/10.1200/jco.2014.60.0734.

[4] a) A. Luetke, P. A. Meyers, I. Lewis, H. Juergens, Cancer Treat Rev 2014, 40 (4), 523, https://doi.org/10.1016/j.ctrv.2013.11.006; b) H. T. Ta, C. R. Dass, I. Larson, P. F. Choong, D. E. Dunstan, Biomaterials 2009, 30 (21), 3605, https://doi.org/10.1016/j.biomaterials.2009.03.022.

[5] N. Omer, M. C. Le Deley, S. Piperno-Neumann, P. Marec-Berard, A. Italiano, N. Corradini, C. Bellera, L. Brugières, N. Gaspar, Eur J Cancer 2017, 75, 98, https://doi.org/10.1016/j.ejca.2017.01.005.

[6] a) A. Lamora, J. Talbot, M. Mullard, B. Brounais-Le Royer, F. Redini, F. Verrecchia, Journal of Clinical Medicine 2016, 5 (11), 96; b) Z. Yao, B. Murali, Q. Ren, X. Luo, D. V. Faget, T. Cole, B. Ricci, D. Thotala, J. Monahan, J. M. van Deursen, D. Baker, R. Faccio, J. K. Schwarz, S. A. Stewart, Cancer Res 2020, 80 (5), 1171, https://doi.org/10.1158/0008-5472.Can-19-2348; c) O. K. Lee, M. J. Coathup, A. E. Goodship, G. W. Blunn, Tissue Eng 2005, 11 (11-12), 1727, https://doi.org/10.1089/ten.2005.11.1727; d) K. C. Stine, E. C. Wahl, L. Liu, R. A. Skinner, J. Vanderschilden, R. C. Bunn, C. O. Montgomery, L. J. Suva, J. Aronson, D. L. Becton, R. W. Nicholas, C. J. Swearingen, C. K. Lumpkin, Jr., Journal of orthopaedic research : official publication of the Orthopaedic Research Society 2014, 32 (3), 464, https://doi.org/10.1002/jor.22527; e) A. Alfranca, L. Martinez-Cruzado, J. Tornin, A. Abarrategi, T. Amaral, E. de Alava, P. Menendez, J. Garcia-Castro, R. Rodriguez, Cellular and Molecular Life Sciences 2015, 72 (16), 3097, https://doi.org/10.1007/s00018-015-1918-y.

[7] F. E. Freeman, R. Burdis, O. R. Mahon, D. J. Kelly, N. Artzi, Advanced Healthcare Materials n/a (n/a), 2101296, https://doi.org/10.1002/adhm.202101296.

[8] J. Friesenbichler, W. Maurer-Ertl, M. Bergovec, L. A. Holzer, K. Ogris, L. Leitner, A. Leithner, Sci Rep-Uk 2017, 7 (1), 1736, https://doi.org/10.1038/s41598-017-02048-w.

[9] a) B. Cai, L. Huang, J. Wang, D. Sun, C. Zhu, Y. Huang, S. Li, Z. Guo, L. Liu, G. Feng, Y. Li, L. Zhang, Bioconjug Chem 2021, 32 (10), 2184, https://doi.org/10.1021/acs.bioconjchem.1c00367; b) W. Diwu, X. Dong, O. Nasif, S. A. Alharbi, J. Zhao, W. Li, Front Cell Dev Biol 2020, 8, 631107, https://doi.org/10.3389/fcell.2020.631107; c) S. Dong, Y. N. Zhang, J. Wan, R. Cui, X. Yu, G. Zhao, K. Lin, J Mater Chem B 2020, 8 (3), 368, https://doi.org/10.1039/c9tb02383f; d) C. He, C. Dong, L. Yu, Y. Chen, Y. Hao, Adv Sci (Weinh) 2021, 8 (19), e2101739, https://doi.org/10.1002/advs.202101739; e) D. Li, W. Nie, L. Chen, D. McCoul, D. Liu, X. Zhang, Y. Ji, B. Yu, C. He, J Biomed Nanotechnol 2018, 14 (12), 2003, https://doi.org/10.1166/jbn.2018.2646; f) J. Liao, K. Shi, Y. Jia, Y. Wu, Z. Qian, Bioact Mater 2021, 6 (8), 2221, https://doi.org/10.1016/j.bioactmat.2021.01.006; g) J. Long, W. Zhang, Y. Chen, B. Teng, B. Liu, H. Li, Z. Yao, D. Wang, L. Li, X. F. Yu, L. Qin, Y. Lai, Biomaterials 2021, 275, 120950, https://doi.org/10.1016/j.biomaterials.2021.120950; h) J. W. Lu, F. Yang, Q. F. Ke, X. T. Xie, Y. P. Guo, Nanomedicine 2018, 14 (3), 811, https://doi.org/10.1016/j.nano.2017.12.025; i) Y. Lu, L. Li, M. Li, Z. Lin, L. Wang, Y. Zhang, Q. Yin, H. Xia, G. Han, Bioconjug Chem 2018, 29 (9), 2982, https://doi.org/10.1021/acs.bioconjchem.8b00400; j) W. E. G. Müller, M. Ackermann, B. Al-Nawas, L. A. R. Righesso, R. Muñoz-Espí, E. Tolba, M. Neufurth, H. C. Schröder, X. Wang, Acta Biomater 2020, 118, 233, https://doi.org/10.1016/j.actbio.2020.10.023; k) L. Wang, Q. Yang, M. Huo, D. Lu, Y. Gao, Y. Chen, H. Xu, Adv Mater 2021, 33 (31), e2100150, https://doi.org/10.1002/adma.202100150; l) C. Yang, H. Ma, Z. Wang, M. R. Younis, C. Liu, C. Wu, Y. Luo, P. Huang, Adv Sci (Weinh) 2021, 8 (20), e2100894, https://doi.org/10.1002/advs.202100894; m) Q. Yang, H. Yin, T. Xu, D. Zhu, J. Yin, Y. Chen, X. Yu, J. Gao, C. Zhang, Y. Chen, Y. Gao, Small 2020, 16 (14), e1906814, https://doi.org/10.1002/smll.201906814; n) M. Yao, Q. Zou, W. Zou, Z. Xie, Z. Li, X. Zhao, C. Du, Biomater Sci 2021, 9 (9), 3319, https://doi.org/10.1039/d0bm01785j; o) J. Yin, S. Pan, X. Guo, Y. Gao, D. Zhu, Q. Yang, J. Gao, C. Zhang, Y. Chen, Nanomicro Lett 2021, 13 (1), 30, https://doi.org/10.1007/s40820-020-00547-6; p) D. Zhang, S. Cheng, J. Tan, J. Xie, Y. Zhang, S. Chen, H. Du, S. Qian, Y. Qiao, F. Peng, X. Liu, Bioact Mater 2022, 17, 394, https://doi.org/10.1016/j.bioactmat.2022.01.032; q) Z. F. Zhou, T. W. Sun, F. Chen, D. Q. Zuo, H. S. Wang, Y. Q. Hua, Z. D. Cai, J. Tan, Biomaterials 2017, 121, 1, https://doi.org/10.1016/j.biomaterials.2016.12.031; r) C. Zhu, M. He, D. Sun, Y. Huang, L. Huang, M. Du, J. Wang, J. Wang, Z. Li, B. Hu, Y. Song, Y. Li, G. Feng, L. Liu, L. Zhang, ACS Appl Mater Interfaces 2021, 13 (40), 47327, https://doi.org/10.1021/acsami.1c10898; s) H. Zhuang, C. Qin, M. Zhang, J. Ma, D. Zhai, B. Ma, N. Ma, Z. Huan, C. Wu, Biofabrication 2021, 13 (4), https://doi.org/10.1088/1758-5090/ac19c7; t) F. Yang, J. Lu, Q. Ke, X. Peng, Y. Guo, X. Xie, Sci Rep 2018, 8 (1), 7345, https://doi.org/10.1038/s41598-018-25595-2.

[10] K. S. Nørregaard, H. J. Jürgensen, H. Gårdsvoll, L. H. Engelholm, N. Behrendt, K. Søe, Int J Mol Sci 2021, 22 (13), https://doi.org/10.3390/ijms22136865.

[11] a) Y. Deng, H. Zhou, P. Gu, X. Fan, Invest Ophthalmol Vis Sci 2014, 55 (9), 6016, https://doi.org/10.1167/iovs.14-14977; b) A. Gilam, J. Conde, D. Weissglas-Volkov, N. Oliva, E. Friedman, N. Artzi, N. Shomron, Nat Commun 2016, 7, 12868, https://doi.org/10.1038/ncomms12868.

[12] B. Yan, Q. Guo, F. J. Fu, Z. Wang, Z. Yin, Y. B. Wei, J. R. Yang, Onco Targets Ther 2015, 8, 539, https://doi.org/10.2147/OTT.S75899.

[13] a) M. J. Schmitt, C. Margue, I. Behrmann, S. Kreis, Curr Mol Med 2013, 13 (4), 572, https://doi.org/10.2174/1566524011313040009; b) S. Sur, R. Steele, X. Shi, R. B. Ray, Cells 2019, 8 (11), https://doi.org/10.3390/cells8111455; c) L. H. Wang, J. Huang, C. R. Wu, L. Y. Huang, J. Cui, Z. Z. Xing, C. Y. Zhao, Mol Med Rep 2018, 17 (2), 2113, https://doi.org/10.3892/mmr.2017.8145; d) G. Song, Y. Zhou, R. Chen, Q. Li, B. Shan, Y. Duan, Y. Wang, Clin Lab 2016, 62 (9), 1739, https://doi.org/10.7754/Clin.Lab.2016.160204; e) Y. Teng, Y. Zhang, K. Qu, X. Yang, J. Fu, W. Chen, X. Li, Oncotarget 2015, 6 (38), 40799, https://doi.org/10.18632/oncotarget.5695.

[14] J. Shin, H. G. Shim, T. Hwang, H. Kim, S.-H. Kang, Y.-S. Dho, S.-H. Park, S. J. Kim, C.-K. Park, Cancer Cell International 2017, 17 (1), 104, https://doi.org/10.1186/s12935-017-0476-9.

[15] a) Z. Li, M. Q. Hassan, M. Jafferji, R. I. Aqeilan, R. Garzon, C. M. Croce, A. J. van Wijnen, J. L. Stein, G. S. Stein, J. B. Lian, J Biol Chem 2009, 284 (23), 15676, https://doi.org/10.1074/jbc.M809787200; b) M. Rossi, M. R. Pitari, N. Amodio, M. T. Di Martino, F. Conforti, E. Leone, C. Botta, F. M. Paolino, T. Del Giudice, E. Iuliano, M. Caraglia, M. Ferrarini, A. Giordano, P. Tagliaferri, P. Tassone, J Cell Physiol 2013, 228 (7), 1506, https://doi.org/10.1002/jcp.24306.

[16] a) H. X. Chen, X. X. Xu, B. Z. Tan, Z. Zhang, X. D. Zhou, Cell Physiol Biochem 2017, 41 (3), 933, https://doi.org/10.1159/000460510; b) J. H. Fang, H. C. Zhou, C. Zeng, J. Yang, Y. Liu, X. Huang, J. P. Zhang, X. Y. Guan, S. M. Zhuang, Hepatology 2011, 54 (5), 1729, https://doi.org/10.1002/hep.24577; c) Y. Li, B. Cai, L. Shen, Y. Dong, Q. Lu, S. Sun, S. Liu, S. Ma, P. X. Ma, J. Chen, Cancer Lett 2017, 397, 111, https://doi.org/10.1016/j.canlet.2017.03.032; S. Sur, R. Steele, X. Shi, R. B. Ray, Cells 2019, 8 (11), 1455, https://doi.org/10.3390/cells8111455.

[17] a) K. Zhu, L. Liu, J. Zhang, Y. Wang, H. Liang, G. Fan, Z. Jiang, C. Y. Zhang, X. Chen, G. Zhou, Protein Cell 2016, 7 (6), 434, https://doi.org/10.1007/s13238-016-0277-2; b) W. Xu, Z. Li, X. Zhu, R. Xu, Y. Xu, Med Sci Monit 2018, 24, 8812, https://doi.org/10.12659/msm.911972; c) K. Zhang, C. Zhang, L. Liu, J. Zhou, Int J Clin Exp Pathol 2014, 7 (9), 5701.

[18] D. J. Luo, L. J. Li, H. F. Huo, X. Q. Liu, H. W. Cui, D. M. Jiang, Eur Rev Med Pharmacol Sci 2019, 23 (4), 1434, https://doi.org/10.26355/eurrev_201902_17100.

[19] Y. Chen, D. Y. Gao, L. Huang, Adv Drug Deliv Rev 2015, 81, 128, https://doi.org/10.1016/j.addr.2014.05.009.

[20] a) M. Guvendiren, H. D. Lu, J. A. Burdick, Soft Matter 2012, 8 (2), 260, https://doi.org/10.1039/C1SM06513K; b) L. L. Wang, Y. Liu, J. J. Chung, T. Wang, A. C. Gaffey, M. Lu, C. A. Cavanaugh, S. Zhou, R. Kanade, P. Atluri, E. E. Morrisey, J. A. Burdick, Nature Biomedical Engineering 2017, 1 (12), 983, https://doi.org/10.1038/s41551-017-0157-y.

[21] E. Larrañeta, M. Henry, N. J. Irwin, J. Trotter, A. A. Perminova, R. F. Donnelly, Carbohydr Polym 2018, 181, 1194, https://doi.org/10.1016/j.carbpol.2017.12.015.

[22] N. Puigmal, P. Dosta, Z. Solhjou, K. Yatim, C. Ramírez, J. Y. Choi, J. B. Alhaddad, A. P. Cosme, J. Azzi, N. Artzi, Advanced Functional Materials 2021, 31 (44), 2100128, https://doi.org/https://doi.org/10.1002/adfm.202100128.

[23] P. Dosta, I. Tamargo, V. Ramos, S. Kumar, D. W. Kang, S. Borrós, H. Jo, Advanced Healthcare Materials 2021, 10 (15), 2001894, https://doi.org/https://doi.org/10.1002/adhm.202001894.

[24] a) J. C. Sunshine, S. B. Sunshine, I. Bhutto, J. T. Handa, J. J. Green, PLOS ONE 2012, 7 (5), e37543, https://doi.org/10.1371/journal.pone.0037543; b) X. Gao, Z. Jin, X. Tan, C. Zhang, C. Zou, W. Zhang, J. Ding, B. C. Das, K. Severinov, I. I. Hitzeroth, Journal of Controlled Release 2020, 321, 654; c) D. M. Lynn, R. Langer, Journal of the American Chemical Society 2000, 122 (44), 10761.

[25] T. Zhang, X. Xue, H. Peng, Mol Ther 2019, 27 (6), 1183, https://doi.org/10.1016/j.ymthe.2019.03.020.

[26] A. Mikai, M. Ono, I. Tosa, H. T. T. Nguyen, E. S. Hara, S. Nosho, A. Kimura-Ono, K. Nawachi, T. Takarada, T. Kuboki, T. Oohashi, Int J Mol Sci 2020, 21 (19), 7028.

[27] P. Dosta, V. Ramos, S. Borrós, Molecular Systems Design & Engineering 2018, 3 (4), 677, https://doi.org/10.1039/C8ME00006A.

[28] P. Dosta, C. Demos, V. Ramos, D. W. Kang, S. Kumar, H. Jo, S. Borrós, Cardiovascular Engineering and Technology 2021, 12 (1), 114, https://doi.org/10.1007/s13239-021-00518-x.

[29] N. Segovia, P. Dosta, A. Cascante, V. Ramos, S. Borrós, Acta Biomaterialia 2014, 10 (5), 2147, https://doi.org/https://doi.org/10.1016/j.actbio.2013.12.054.

[30] B. T. Grisez, J. J. Ray, P. A. Bostian, J. E. Markel, B. A. Lindsey, J Orthop Res 2018, https://doi.org/10.1002/jor.23868.

[31] M. Cortini, N. Baldini, S. Avnet, Front Physiol 2019, 10, 814, https://doi.org/10.3389/fphys.2019.00814.

[32] a) J. Han, B. Yong, C. Luo, P. Tan, T. Peng, J. Shen, World Journal of Surgical Oncology 2012, 10 (1), 37, https://doi.org/10.1186/1477-7819-10-37; b) J. Zhou, T. Liu, W. Wang, Medicine 2018, 97 (44), e13051, https://doi.org/10.1097/MD.0000000000013051; c) P. Kunz, H. Sähr, B. Lehner, C. Fischer, E. Seebach, J. Fellenberg, BMC Cancer 2016, 16 (1), 223, https://doi.org/10.1186/s12885-016-2266-5; d) J. Wang, Q. Shi, T.-x. Yuan, Q.-l. Song, Y. Zhang, Q. Wei, L. Zhou, J. Luo, G. Zuo, M. Tang, T.-C. He, Y. Weng, Clinica Chimica Acta 2014, 433, 225, https://doi.org/https://doi.org/10.1016/j.cca.2014.03.023.

[33] a) G. Hulsart-Billström, K. Bergman, B. Andersson, J. Hilborn, S. Larsson, K. B. Jonsson, J Tissue Eng Regen Med 2015, 9 (7), 799, https://doi.org/10.1002/term.1655; b) S. W. Jung, J. H. Byun, S. H. Oh, T. H. Kim, J. S. Park, G. J. Rho, J. H. Lee, Carbohydr Polym 2018, 180, 216, https://doi.org/10.1016/j.carbpol.2017.10.029; c) C. Martínez-Álvarez, B. González-Meli, B. Berenguer-Froehner, I. Paradas-Lara, Y. López-Gordillo, C. Rodríguez-Bobada, P. González, M. Chamorro, P. Arias, J. Hilborn, I. Casado-Gómez, E. Martínez-Sanz, J Surg Res 2013, 183 (2), 654, https://doi.org/10.1016/j.jss.2013.03.009; d) E. Martínez-Sanz, D. A. Ossipov, J. Hilborn, S. Larsson, K. B. Jonsson, O. P. Varghese, J Control Release 2011, 152 (2), 232, https://doi.org/10.1016/j.jconrel.2011.02.003; e) E. Martínez-Sanz, O. P. Varghese, M. Kisiel, T. Engstrand, K. M. Reich, M. Bohner, K. B. Jonsson, T. Kohler, R. Müller, D. A. Ossipov, J. Hilborn, J Tissue Eng Regen Med 2012, 6 Suppl 3, s15, https://doi.org/10.1002/term.1593; f) S. H. Park, J. Y. Park, Y. B. Ji, H. J. Ju, B. H. Min, M. S. Kim, Acta Biomater 2020, 117, 108, https://doi.org/10.1016/j.actbio.2020.09.013.

[34] P. D. Ottewell, J. K. Woodward, D. V. Lefley, C. A. Evans, R. E. Coleman, I. Holen, Mol Cancer Ther 2009, 8 (10), 2821, https://doi.org/10.1158/1535-7163.MCT-09-0462.

[35] a) F. E. Freeman, P. Pitacco, L. H. A. van Dommelen, J. Nulty, D. C. Browe, J. Y. Shin, E. Alsberg, D. J. Kelly, Sci Adv 2020, 6 (33), eabb5093, https://doi.org/10.1126/sciadv.abb5093; b) Y. M. Kolambkar, K. M. Dupont, J. D. Boerckel, N. Huebsch, D. J. Mooney, D. W. Hutmacher, R. E. Guldberg, Biomaterials 2011, 32 (1), 65, https://doi.org/https://doi.org/10.1016/j.biomaterials.2010.08.074; c) J. D. Boerckel, Y. M. Kolambkar, K. M. Dupont, B. A. Uhrig, E. A. Phelps, H. Y. Stevens, A. J. García, R. E. Guldberg, Biomaterials 2011, 32 (22), 5241, https://doi.org/https://doi.org/10.1016/j.biomaterials.2011.03.063; d) M. Kisiel, M. M. Martino, M. Ventura, J. A. Hubbell, J. Hilborn, D. A. Ossipov, Biomaterials 2013, 34 (3), 704, https://doi.org/https://doi.org/10.1016/j.biomaterials.2012.10.015.

[36] A. Gupta, R. Bahal, M. Gupta, P. M. Glazer, W. M. Saltzman, Journal of controlled release : official journal of the Controlled Release Society 2016, 240, 302, https://doi.org/10.1016/j.jconrel.2016.01.005.

[37] P. Dosta, N. Segovia, A. Cascante, V. Ramos, S. Borrós, Acta Biomaterialia 2015, 20, 82, https://doi.org/https://doi.org/10.1016/j.actbio.2015.03.029.

[38] A. Tchoryk, V. Taresco, R. H. Argent, M. Ashford, P. R. Gellert, S. Stolnik, A. Grabowska, M. C. Garnett, Bioconjugate Chemistry 2019, 30 (5), 1371, https://doi.org/10.1021/acs.bioconjchem.9b00136.

[39] a) M. Vinci, S. Gowan, F. Boxall, L. Patterson, M. Zimmermann, W. Court, C. Lomas, M. Mendiola, D. Hardisson, S. A. Eccles, BMC Biology 2012, 10 (1), 29, https://doi.org/10.1186/1741-7007-10-29; b) A. Ivascu, M. Kubbies, J Biomol Screen 2006, 11 (8), 922, https://doi.org/10.1177/1087057106292763; c) A. S. Mikhail, S. Eetezadi, C. Allen, PLoS One 2013, 8 (4), e62630, https://doi.org/10.1371/journal.pone.0062630.

[40] a) A. Pluen, Y. Boucher, S. Ramanujan, T. D. McKee, T. Gohongi, E. d. Tomaso, E. B. Brown, Y. Izumi, R. B. Campbell, D. A. Berk, R. K. Jain, Proceedings of the National Academy of Sciences 2001, 98 (8), 4628, https://doi.org/doi:10.1073/pnas.081626898; b) H. Cabral, Y. Matsumoto, K. Mizuno, Q. Chen, M. Murakami, M. Kimura, Y. Terada, M. R. Kano, K. Miyazono, M. Uesaka, N. Nishiyama, K. Kataoka, Nature Nanotechnology 2011, 6 (12), 815, https://doi.org/10.1038/nnano.2011.166.

[41] H.-X. Wang, Z.-Q. Zuo, J.-Z. Du, Y.-C. Wang, R. Sun, Z.-T. Cao, X.-D. Ye, J.-L. Wang, K. W. Leong, J. Wang, Nano Today 2016, 11 (2), 133, https://doi.org/https://doi.org/10.1016/j.nantod.2016.04.008.

[42] a) K. B. Jones, Z. Salah, S. Del Mare, M. Galasso, E. Gaudio, G. J. Nuovo, F. Lovat, K. LeBlanc, J. Palatini, R. L. Randall, S. Volinia, G. S. Stein, C. M. Croce, J. B. Lian, R. I. Aqeilan, Cancer Res 2012, 72 (7), 1865, https://doi.org/10.1158/0008-5472.CAN-11-2663; b) N. Dai, Z. Y. Zhong, Y. P. Cun, Y. Qing, C. Chen, P. Jiang, M. X. Li, D. Wang, Neoplasma 2013, 60 (4), 384, https://doi.org/10.4149/neo_2013_050.

[43] a) T. Nakasa, M. Yoshizuka, M. Andry Usman, E. Elbadry Mahmoud, M. Ochi, Curr Genomics 2015, 16 (6), 441, https://doi.org/10.2174/1389202916666150817213630; b) K. Kapinas, C. Kessler, T. Ricks, G. Gronowicz, A. M. Delany, The Journal of biological chemistry 2010, 285 (33), 25221, https://doi.org/10.1074/jbc.M110.116137; c) Q. Zeng, Y. Wang, J. Gao, Z. Yan, Z. Li, X. Zou, Y. Li, J. Wang, Y. Guo, Cell Mol Biol Lett 2019, 24, 11, https://doi.org/10.1186/s11658-019-0136-2; d) T. Xia, S. Dong, J. Tian, Int J Mol Med 2020, 46 (2), 709, https://doi.org/10.3892/ijmm.2020.4615.

[44] R. D. Roberts, M. M. Lizardo, D. R. Reed, P. Hingorani, J. Glover, W. Allen-Rhoades, T. Fan, C. Khanna, E. A. Sweet-Cordero, T. Cash, M. W. Bishop, M. Hegde, A. R. Sertil, C. Koelsche, L. Mirabello, D. Malkin, P. H. Sorensen, P. S. Meltzer, K. A. Janeway, R. Gorlick, B. D. Crompton, Cancer 2019, 125 (20), 3514, https://doi.org/10.1002/cncr.32351.

[45] B. Tan, Y. Wu, Y. Wu, K. Shi, R. Han, Y. Li, Z. Qian, J. Liao, ACS Appl Mater Interfaces 2021, 13 (27), 31542, https://doi.org/10.1021/acsami.1c08775.

[46] C. Bailly, X. Thuru, B. Quesnel, NAR Cancer 2020, 2 (1), https://doi.org/10.1093/narcan/zcaa002.

[47] a) P. Dosta, A. M. Cryer, M. Prado, M. Z. Dion, S. Ferber, S. Kalash, N. Artzi, Advanced NanoBiomed Research 2021, 1 (7), 2100006, https://doi.org/https://doi.org/10.1002/anbr.202100006; b) P. Dosta, Cryer, A.M., Dion, M.Z., Shiraishi, T., Langston, S.P., Lok, D., Wang, J., Harrison, S., Ferber, S., Kalash, S., Prado, M., Rodríguez, A.L., Abu-Yousif, A.O., Artzi, N., Research Square 2022, https://doi.org/doi:10.21203/rs.3.rs-1647238/v1.

[48] G. R. Mundy, Nature Reviews Cancer 2002, 2 (8), 584, https://doi.org/10.1038/nrc867.

[49] a) F. E. Freeman, M. Haugh, L. McNamara, Journal of tissue engineering and regenerative medicine 2013, https://doi.org/10.1002/term.1793; b) F. E. Freeman, M. G. Haugh, L. McNamara, Tissue Engineering Part A 2015, 21 (7-8), 1320, https://doi.org/10.1089/ten.tea.2014.0249; c) F. E. Freeman, H. Stevens, P. Owens, R. Guldberg, L. McNamara, Tissue Eng Part A 2016, 22 (19-20), https://doi.org/10.1089/ten.TEA.2015.0339.

[50] F. E. Freeman, D. J. Kelly, Sci Rep 2017, 7 (1), 17042, https://doi.org/10.1038/s41598-017-17286-1.

[51] E. B. Dolan, M. G. Haugh, D. Tallon, C. Casey, L. M. McNamara, J R Soc Interface 2012, 9 (77), 3503, https://doi.org/10.1098/rsif.2012.0520.

[52] F. E. Freeman, D. C. Browe, J. Nulty, S. Von Euw, W. L. Grayson, D. J. Kelly, Eur Cell Mater 2019, 38, 168, https://doi.org/10.22203/eCM.v038a12.

[53] J. Nulty, R. Burdis, D. J. Kelly, Frontiers in Bioengineering and Biotechnology 2021, 9 (469), https://doi.org/10.3389/fbioe.2021.661989.

[54] Y. Zhang, P. Dosta, J. Conde, N. Oliva, M. Wang, N. Artzi, Advanced Healthcare Materials 2020, 9 (4), 1901101, https://doi.org/https://doi.org/10.1002/adhm.201901101.

[55] R. Núñez-Toldrà, P. Dosta, S. Montori, V. Ramos, M. Atari, S. Borrós, Acta Biomaterialia 2017, 53, 152, https://doi.org/https://doi.org/10.1016/j.actbio.2017.01.077.

